# Tree-based Correlation Screen and Visualization for Exploring Phenotype-Cell Type Association in Multiple Sample Single-Cell RNA-Sequencing Experiments

**DOI:** 10.1101/2021.10.27.466024

**Authors:** Boyang Zhang, Zhicheng Ji, Hongkai Ji

## Abstract

Single-cell RNA-seq experiments with multiple samples are increasingly used to discover cell types and their molecular features that may influence samples’ phenotype (e.g. disease). However, analyzing and visualizing the complex cell type-phenotype association remains nontrivial. TreeCorTreat is an open source R package that tackles this problem by using a <u>tree</u>-based <u>cor</u>relation screen to analyze and visualize the association between phenotype and <u>tr</u>anscriptomic f<u>e</u>atures <u>a</u>nd cell <u>t</u>ypes at multiple cell type resolution levels. With TreeCorTreat, one can conveniently explore and compare different feature types, phenotypic traits, analysis protocols and datasets, and evaluate the impacts of potential confounders.

## Background

Single-cell RNA-sequencing (scRNA-seq) has been rapidly transforming biomedical research [1–4]. With the ability to profile the transcriptomes of individual cells, this technology is now widely used to discover new cell types, analyze cell-cell interactions, and map spatial and temporal transcriptional programs [4–7]. Crucial to these applications are computational tools that retrieve, synthesize, and visualize information from the complex and noisy data. In the past few years, many computational methods and tools have been developed to support different scRNA-seq data analysis tasks such as data preprocessing[8–10], normalization[11–15], integration[16–19], imputation[20–25], visualization[26,27], cell clustering[28,29], trajectory inference[30–34], and spatial mapping[6,7,35,36]. A main focus of these tools is to deal with cell-level heterogeneity or analyze the relationship among cells, such as how cells cluster, interact, or how they change progressively along time or space. However, today’s scRNA-seq experiments increasingly generate data from multiple biological samples with different phenotypes, introducing sample-level heterogeneity in addition to the cell-level heterogeneity. For example, since 2020, over 600 COVID-19 patients have been profiled using scRNA-seq by multiple research groups to investigate transcriptomic features associated with the progression of disease severity[37–42]. Similarly, multiple studies have used scRNA-seq to analyze tumor infiltrating lymphocytes (TILs) from immunotherapy-treated patients to search for molecular features associated with treatment response[43–45]. The sample-level variability introduces a new layer of complexity and new analytical needs not met by existing computational tools.

For a scRNA-seq dataset with multiple biological samples and multiple cell types in each sample, an important question is how different cell types are associated with samples’ phenotype such as COVID disease severity or patient’s response to cancer immunotherapy. Answering this question is nontrivial for several reasons. First, a cell type can be associated with phenotype in different ways. Unknown a priori, a phenotype may be associated with either cell type proportion or gene expression or both. Second, cell types in a sample may be defined at multiple resolution levels. For instance, cells in a peripheral blood mononuclear cell (PBMC) sample can be broadly classified into lymphocytes and monocytes. Lymphocytes can be further categorized into B cells, T cells, and natural killer (NK) cells. At a finer resolution, T cells can be further grouped into CD4+ T cells, CD8+ T cells, etc. A priori, it is unknown which cell type resolution level is most appropriate for analyzing the association with the phenotype. Exhaustively analyzing the phenotype-cell type association at all resolutions can easily produce large amounts of results (e.g. tens of thousands of statistically significant features such as differentially expressed genes) that will overwhelm an investigator. How to digest the information to distill key findings remains an important problem that still lacks software support. Third, a sample may have multiple phenotypic traits. Comparing the phenotype-cell type association across different traits or analyzing traits jointly can introduce a new layer of complexity which amplifies the burden of the analysis. Similarly, one trait may be analyzed in multiple datasets. Comparing datasets to examine whether the phenotype-cell type association is consistent across studies can also make the analysis more complex. Last but not least, the association between phenotype and cell proportion or gene expression can be confounded by variables not of primary interest, such as experimental batches. In order to accurately study the phenotype-cell type association, it is important to explore and understand the impacts of potential confounders.

Given all these complexities, exploratory data analysis (EDA) plays an important role in investigating phenotype-cell type association in multi-sample scRNA-seq data. The goal of such an analysis is to learn (1) which cell types and feature types (i.e., cell proportion or gene expression) are associated with and most relevant for studying the phenotype of interest, (2) at which cell type resolution levels does the association exist, (3) how the association varies across different phenotypic traits, datasets, or analysis protocols, and (4) how potential confounders can influence the association. This exploration can help investigators to obtain big pictures of the complex data, which in turn can serve as a foundation for designing and conducting in-depth downstream computational and experimental analyses. For example, once the most relevant cell types are identified, one can then perform more focused analyses to carefully examine features (e.g. differentially expressed genes) of these cell types to develop mechanistic hypotheses and design subsequent experiments to study their functions.

During the exploration, visualization plays an indispensable role in presenting the big picture and key information. Visualization tools that effectively summarize and display information can have a long-lasting impact. Two such examples are heatmap for visualizing gene expression data [46] and sequence logo for visualizing DNA or protein sequence motifs [47,48]. For analyzing multi-sample scRNA-seq data, there is a growing need for a tool that supports convenient visualization of multi-resolution phenotype-cell type associations with the flexibility to handle different feature types, multivariate phenotype, multiple datasets, and potential confounders.

The growing needs for exploring and visualizing phenotype-cell type association in multi-sample scRNA-seq data cannot be readily met by existing computational tools. Tools for analyzing differential expression (DE) may be used to detect phenotype-cell type association in terms of gene expression. However, existing scRNA-seq DE analysis tools, such as Seurat (Wilcoxon test, Student’s T test, Generalized linear model) [17], MAST[49], SCDE[50], scDD[51] and DEsingle[52], mainly address the problem of comparing two cell populations. While they consider cell-level variability, they do not consider sample-level variability in multi-sample studies. DE analysis tools developed for bulk gene expression data, such as limma [53], DESeq [54,55] and edgeR [56], consider sample-level variability and can handle multiple samples. However, they do not consider the fact that a bulk sample may contain multiple cell subpopulations and cannot automatically identify cell subpopulations and analyze DE in each subpopulation. None of the single cell or bulk DE analysis tools above support analysis at multiple cell type resolution levels. Moreover, none of them can support comprehensive exploration of different feature types (e.g. gene expression and cell type proportion) and different types of phenotypic traits (e.g. univariate and multivariate phenotype). They also do not support visualization and cannot be conveniently used to compare results from different datasets, phenotypic traits, and analysis protocols (e.g., before and after adjusting for a potential confounder) (**Additional file 1: Table S1**). One recent method considers cell type hierarchical structure to evaluate cell clustering results in scRNA-seq data, but it does not tackle the problem of identifying phenotype-cell type association in multi-sample studies [57]. For visualization, although tools such as UMAP[26] and tSNE [27] are available to visualize cell-level information, a method that supports convenient visualization of multi-resolution cell type-phenotype association is still lacking.

In order to fill the critical gap for exploring and visualizing phenotype-cell type association in multi-sample scRNA-seq data, we introduce <u>tree</u>-based <u>cor</u>relation screen for phenotype-associated <u>tr</u>anscriptomic f<u>e</u>atures <u>a</u>nd cell <u>t</u>ypes (TreeCorTreat), a software tool that supports multi-resolution and multi-feature type association analysis in single-cell data. We also introduce TreeCorTreat plot as a new visualization tool to summarize the association information in such data. TreeCorTreat and its plotting functions are implemented as an open source R package freely available at https://github.com/byzhang23/TreeCorTreat. Together, they can help users to more efficiently explore, visualize and compare the association analysis results obtained at different cell type resolutions, for different phenotypic traits, and from different datasets or analysis protocols.

## Results

### Tree-based correlation screen for phenotype-cell type association

The objective of TreeCorTreat is to support the exploration and visualization of the multi-resolution cell type-phenotype association in multi-sample scRNA-seq data. TreeCorTreat takes a gene expression matrix (raw count), cell-level metadata and sample-level metadata as input. It provides a whole pipeline to integrate data across samples, identify cell clusters and their hierarchical structure, evaluate the association between sample phenotype and cell type at different resolution levels in terms of both cell type proportion and gene expression, and summarize and visualize the results in a tree structured TreeCorTreat plot. This pipeline consists of six functional modules (**Fig. 1**). The modular structure provides users with the flexibility to skip certain analysis steps and replace them by users’ own data or analysis functions.

**Figure 1.**
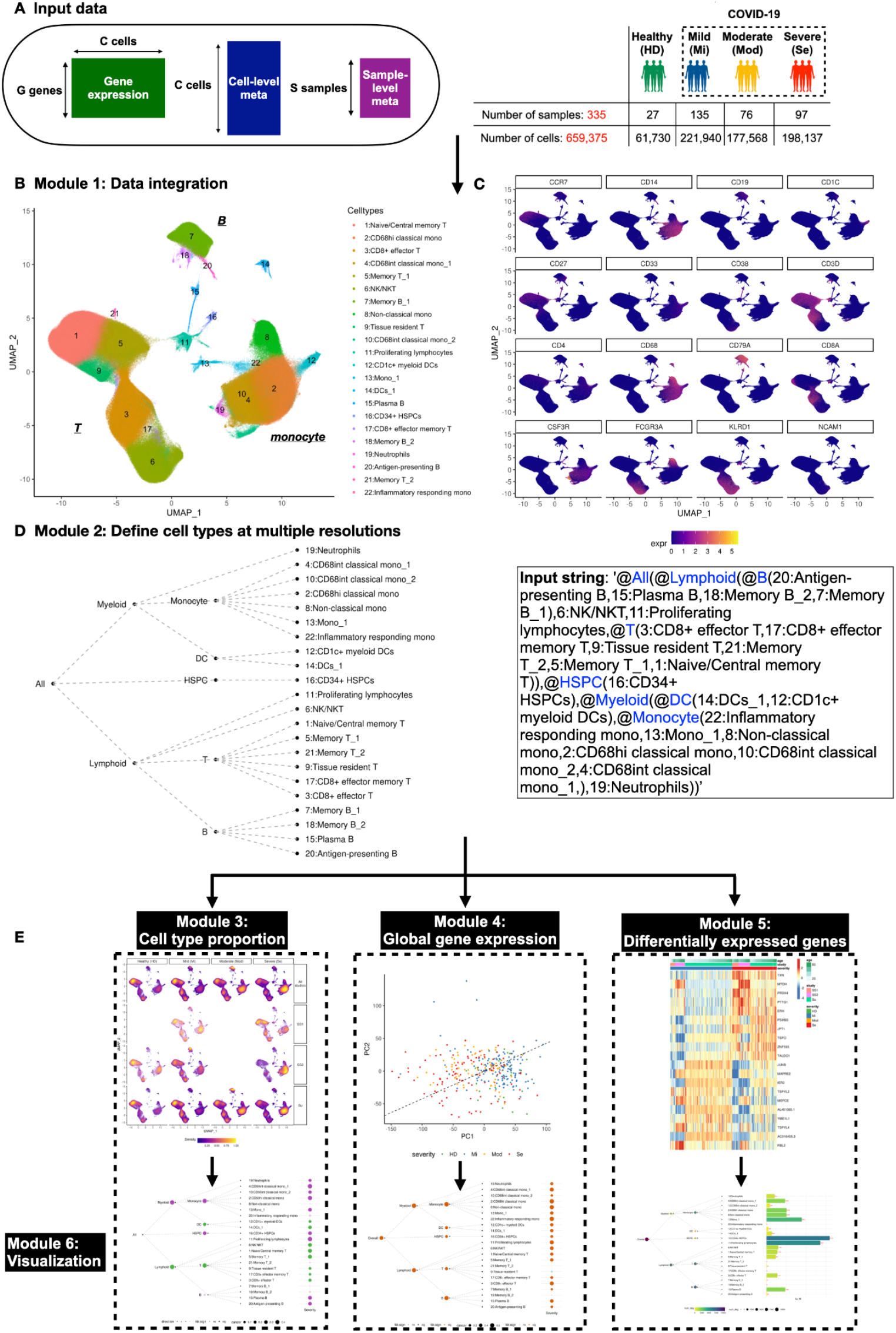
Overview of TreeCorTreat, a tree-based correlation screen and visualization method for exploring multi-sample phenotype-cell type association, using COVID-19 data as a demonstration. (A) TreeCorTreat first takes in a gene expression matrix, cell- and sample-level metadata as input. In the COVID-19 example, a total of 335 samples across 4 disease categories are retained. (B) Using harmony integration and louvain clustering, TreeCorTreat first integrates samples and groups cells into clusters (Module 1). Inferred cell types can be annotated based on cluster-specific differentially expressed genes or gene markers. (C) Expression of 16 known gene markers defining broader cell categories in the COVID-19 samples. (D) Based on prior knowledge, TreeCorTreat can define cell types at multiple resolutions and visualize in a tree structure by parsing an input string (Module 2). (E) TreecorTreat can further analyze the association between samples’ phenotype and cell type proportion (Module 3) and global gene expression of each cell type (Module 4), and detect differentially expressed genes for each cell type (Module 5). The analysis is performed at different cell type resolution levels guided by the tree, and the results can be summarized and visualized in a TreeCorTreat plot (Module 6).

Below we first demonstrate the basic TreeCorTreat pipeline using an analysis of PBMC scRNA-seq data from COVID-19 patients. Here a biological question of interest is how different cell types are associated with COVID-19 disease severity. In particular, we are interested in knowing which cell types are most relevant for studying disease severity and whether the disease severity is linked to changes in cell type composition or gene expression or both.

We downloaded publicly available scRNA-seq data from 371 PBMC samples generated by two studies, Su et al. [37] and Schulte-Schrepping et al. [38]. The samples in Su et al. were processed using a droplet-based single-cell platform (10x Chromium). The samples from Schulte-Schrepping et al. represent two different patient cohorts. The first cohort (SS1) were processed using a droplet-based single-cell platform (10x Chromium), and the second cohort (SS2) were processed using a microwell-based scRNA-seq system (BD Rhapsody). Thus, there were three patient cohorts (Su, SS1, SS2) consisting of 371 samples and 747,697 cells in total. After preprocessing and filtering out low quality cells and samples, 659,375 cells from 335 samples were retained for subsequent analyses (**Additional file 5: Fig. S1A, Methods**). These 335 samples consist of 27 samples from Healthy Donors (HD), 135 Mild (Mi), 76 Moderate (Mod) and 97 Severe (Se) COVID-19 disease samples (**Fig. 1A**).

#### Step 1 - Data integration (Module 1)

The analysis begins with integrating data across samples. TreeCorTreat’s data integration pipeline is wrapped into a function ‘*treecor_harmony()*’. This pipeline uses Harmony [16] as its default data integration algorithm. Using this pipeline, data from different COVID-19 samples and cohorts were integrated and embedded into a common low-dimensional space. Cells from the three cohorts (Su, SS1, SS2) were mixed well in the low-dimensional embedding (**Additional file 5: Fig. S1B**). Unsupervised louvain clustering using the harmonized data identified 22 cell clusters (**Fig. 1B**). The pipeline also identifies genes differentially expressed across clusters. Based on the top differentially expressed genes and known cell type marker genes, we annotated the cell type of each cell cluster by manually assigning a name (**Fig. 1B,C, Additional file 5: Fig. S2**), which are consistent with previous findings [39]. Users can choose to skip this integration step (i.e. Module 1) by providing their own integrated data, such as those obtained from Seurat [17], along with cells’ clustering results through the input gene expression matrix and cell-level metadata.

#### Step 2 - Define cell types at multiple resolutions (Module 2)

To support analyses at different cell type granularity levels, the initial cell clusters from step 1 are hierarchically merged into coarse-grained clusters. The merging process is guided by a tree (dendrogram). The nodes in the tree represent cell types defined at different resolution levels, with the leaf nodes corresponding to the highest resolution. The tree can be either derived from the data using hierarchical clustering (data-driven) or specified by users based on their prior knowledge (knowledge-based). The data-driven approach is wrapped into a function ‘*extract_hrchy_seurat()*’. In the knowledge-based approach, users provide a string to describe the parent-children relationship of clusters at different granularity levels. The string will be parsed by a function ‘*extract_hrchy_string()*’ to create the tree. For example, the string ‘@White_blood_cells(@Lymphocytes(T_cells,B_cells),Monocytes,Granulocytes)’ means that white blood cells can be categorized into lymphocytes, monocytes and granulocytes, and lymphocytes can be further categorized into T cells and B cells. To demonstrate, we constructed a tree for the COVID-19 data based on prior knowledge (**Fig. 1D**). Below we use this tree to demonstrate the downstream analyses.

For each tree node (i.e. cell type), TreeCorTreat can analyze its association with sample phenotype in three different ways: (a) association with cell type proportion, (b) association with global gene expression, and (c) phenotype-associated differentially expressed genes (DEGs) (**Fig. 1E**).

#### Step 3.a - Identify association between cell type proportion and sample phenotype (Module 3)

The analysis of cell type proportion is wrapped into a function ‘*treecor_ctprop()*’. For each leaf node, we compute the proportion of cells in that node in each sample and then report the Pearson correlation (default) or Spearman’s correlation (optional) between the cell type proportion and sample phenotype. For an intermediate node, the calculation is more complicated and can be done in multiple ways which will be discussed in detail later. By default, we use a simple approach (referred to as ‘Aggregate’) that treats all cells from its descendant nodes as a combined cell cluster and computes the proportion of cells in the combined cluster. The correlation between the cell type proportion and phenotype is then computed. The statistical significance of the correlation is determined using permutation test p-values after adjusting for multiple testing across all tree nodes using Benjamini & Yekutieli (BY) procedure [58]. To demonstrate, we applied the default procedure to COVID-19 samples from the Su dataset. The analysis of the other two sample cohorts (SS1 and SS2) will be discussed later. We coded disease severity as a numerical variable (HD = 1, Mi = 2, Mod = 3, Se = 4). We found that the proportion of lymphoid cells in PBMC, particularly the proportion of T cells, is negatively correlated with disease severity (**Fig. 2A,B**), consistent with previous reports that lymphopenia (i.e. reduced level of lymphocytes) is common in severe COVID-19 patients [59,60]. While the overall proportion of lymphoid cells negatively correlates with the severity, several lymphoid subpopulations such as proliferating lymphocytes and B cells showed positive correlation with the severity, indicating heterogeneous responses to the virus infection among different lymphoid cell subtypes. Among T cells, the proportion of naive and memory T cells and CD8+ effector T cells showed stronger correlation with severity than the other T cell subclusters. While most T cell subclusters (i.e. leaf nodes) showed negative correlation with severity, the negative correlation was stronger at the aggregated T cell population level (i.e. the intermediate node labeled as ‘T’) (**Fig. 2B**, **Additional file 5: Fig. S3A**), suggesting that the abundance of the aggregated T cell population in PBMC may serve as a better clinical variable for monitoring COVID-19 disease severity than the abundance of each individual T cell subset. Unlike lymphoid cells, we found that the proportion of myeloid cells in PBMC is positively correlated with disease severity. Among the myeloid cells, monocytes, particularly CD68+ classical monocytes with intermediate or high CD68 expression, showed positive correlation with severity, whereas dendritic cells (DC) showed negative correlation with severity. Overall, the strongest correlates with disease severity in terms of cell type proportion are the abundance of myeloid/monocyte/CD68+ classical monocytes (positive) and lymphoid/T cells (negative) (**Fig. 2B**). Besides using Pearson or Spearman’s correlation to describe association, users also have an option to fit a regression model for each node using cell type proportion as the response variable and sample phenotype as the explanatory variable. The association can then be characterized using a likelihood ratio test between two nested models with and without including the phenotype in the explanatory variables. The test will report the likelihood ratio statistics and multiple testing (BY) corrected p-values. The Pearson and Spearman’s correlation can handle sample phenotypes that are binary or continuous variables or take ordered categorical values, but they cannot handle a sample phenotype with multiple unordered categories. The regression approach, on the other hand, can handle all these data types.

**Figure 2.**
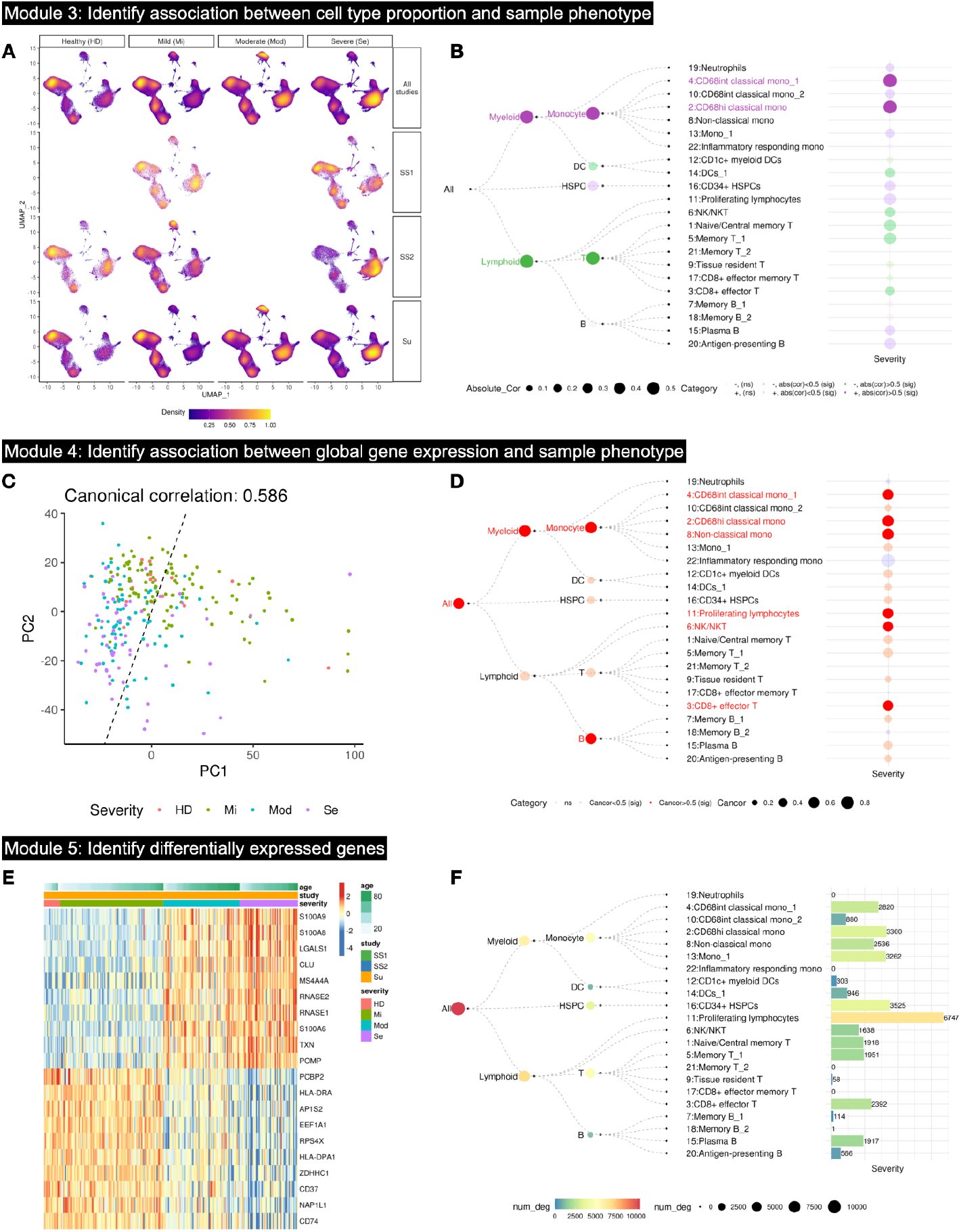
TreeCorTreat analyses of a single COVID-19 dataset (Su et al.). (A)-(B): Module 3 identifies association between cell type proportion and disease severity. (A) TreeCorTreat provides a global view of disease severity’s localization via a density plot of all cells on the UMAP, stratified by disease severity (Healthy (HD), Mild (Mi), Moderate (Mod) and Severe (Se)) and cohort (all studies, SS1, SS2 and Su). (B) Using default setting (aggregate), TreeCorTreat calculates the Pearson correlation between cell type proportion and disease severity for each tree node. In the balloon plot, circle size represents the magnitude of absolute value of Pearson correlation. Colors indicate six categories defined by adjusted p-value (‘sig’ or ‘ns’) and effect size (‘abs(cor) > 0.5’ or ‘abs(cor) ≤ 0.5’), with green for negative correlation and purple for positive correlation. ‘sig’ represents statistically significant after multiple testing adjustments (i.e., adjusted p-value ≤ 0.05) and ‘ns’ represents non-significant. ‘abs(cor)’ stands for absolute value of Pearson correlation. Colored labels highlight statistically significant cell types with a relatively large effect size. For the root node ‘All’ and cell cluster ‘21:Memory T_2’, there is no correlation calculated because the cell type proportions are aggregated as 1 for all samples at root node and cluster 21 contains cells only from 1 sample. (C)-(D): Module 4 identifies association between global gene expression and sample phenotype. (C) PCA plot of sample-level pseudobulk in ‘4:CD48int classical mono_1’ cell type for individual samples with color annotating disease severity. (D) TreeCorTreat plot of association between disease severity and global gene expression of each cell type. Circle size represents the magnitude of canonical correlation and color indicates three categories defined by multiple testing adjusted p-value and effect size (blue for non-significant, light red for significant with canonical correlation<0.5 and solid red for significant with canonical correlation>0.5). Red colored labels represent statistically significant (adjusted p-value ≤ 0.05) cell types with a relatively large effect size (canonical correlation>0.5). Due to a small sample size (n=6) for cell cluster 22:Inflammatory responding monocytes, the canonical correlation is not statistically significant even though its value is large. (E)-(F): Module 5 conducts differential expression analysis. (E) Heatmap of top 10 positively and top 10 negatively differential expressed genes associated with disease severity in ‘4:CD48int classical mono_1’ cell type. Each row is a differentially expressed gene and each column is a sample ordered by disease severity group and age. (F) TreeCorTreat plot of number of differentially expressed genes (DEGs) for each cell type, using adjusted p-value≤0.05 as a cutoff. Color, circle size and bar height represent the number of DEGs.

#### Step 3.b - Identify association between global gene expression and sample phenotype (Module 4)

The goal of this module is to evaluate whether the sample phenotype is associated with the global gene expression profile of each cell type. It is implemented as a function ‘*treecor_expr()*’. Depending on how intermediate tree nodes are handled, the analysis can be run in several different modes. In the default mode, we aggregate cells in each tree node into a pseudobulk gene expression profile per sample. For each tree node, the pseudobulk profiles are used to embed samples into a low-dimensional principal component (PC) space. In this low-dimensional embedding, samples with similar gene expression profiles are located close to each other (**Fig. 2C**). For sample phenotypes that are binary, continuous, or ordered categorical variables, the canonical correlation analysis (CCA) is then used to compute the canonical correlation between the phenotype and samples’ locations in the low-dimensional embedding. The statistical significance of the canonical correlation is characterized using permutation test p-values after multiple testing correction. To demonstrate, we applied this default procedure to samples from the Su dataset. As one example, **Fig. 2C** shows the low-dimensional embedding of samples based on the gene expression profile of cluster 4, which contains classical monocytes with intermediate CD68 levels (i.e. cluster ‘4:CD68int classical mono_1’). One can see that samples’ disease status progressively changes from less severe (healthy and mild) to more severe (moderate and severe) along the direction indicated by the dashed line which corresponds to the first canonical correlation component found by CCA (i.e. the direction that maximizes the correlation between the severity and samples’ coordinates in the embedded space). **Fig. 2D** and **Additional file 5: Fig. S3B** summarize the results for all cell clusters. We found that gene expression profiles of myeloid/monocytes (incl. CD68+ classical monocytes and non-classical monocytes), proliferating lymphocytes, NK/NKT cells, CD8+ effector T cells, and B cells (intermediate node) are most strongly associated with disease severity with canonical correlation > 0.5 (dark red balloons). Many other cell types also showed statistically significant association with severity, but with lower canonical correlation (light red balloons). Of note, cell types associated with phenotype in terms of cell type proportion and cell types associated with phenotype in terms of gene expression are not necessarily the same. For example, non-classical monocytes (cluster ‘8:Non-classical mono’) showed almost no correlation with severity in terms of cell type proportion, but it was strongly correlated with severity in terms of global gene expression. For a sample phenotype with multiple unordered categories, the association between the phenotype and samples’ locations in the low-dimensional embedding will be characterized using a Multivariate Analysis of Variance (MANOVA, Pillai-Bartlett) type analysis (a special case of regression) which computes a ratio of between-group variance versus within-group variance for each node using samples’ embedded locations. Statistical significance is characterized using permutation p-values after multiple testing (BY) correction.

#### Step 3.c - Identify differentially expressed genes (Module 5)

This module, which is wrapped into a function ‘*treecor_deg()*’, will identify differentially expressed genes (DEGs). For each tree node, this function aggregates cells in that node into a pseudobulk gene expression profile per sample. The limma[53] model is then applied to detect DEGs associated with the sample phenotype. The DEGs are exported into csv files, and the results are summarized by visualizing the number of differential genes for each tree node (**Fig. 2F**). For example, when treating disease severity as a numerical variable (HD = 1, Mi = 2, Mod =3, Se = 4) in the Su et al., proliferating lymphocytes showed the largest number of DEGs. Other cell types with a large number of DEGs include CD68+ classical monocytes and non-classical monocytes, CD34+ hematopoietic stem and progenitor cells (HSPCs), NK/NKT cells, naive and a subset of memory T cells, CD8+ effector T cells, and plasma B cells. By contrast, much fewer DEGs were detected in neutrophils, inflammatory responding monocytes, tissue resident T cells, CD8+ effector memory T cells, and memory B cells. **Fig. 2E** shows the top DEGs in the ‘4:CD68int classical mono_1’ cell cluster. Interestingly, three genes of the S100A gene family had elevated gene expression in moderate and severe samples compared to healthy and mild samples, consistent with the fact that moderate/severe COVID-19 patients have a strong systemic inflammatory effect[40].

#### Step 4 - Visualization (Module 6)

The TreeCorTreat package provides a series of functions to visualize both the intermediate and final results. For example, the ‘*treecor_celldensityplot()*’ function shows cell type proportions for different sample groups (**Fig. 2A**). The ‘*treecor_samplepcaplot()*’ function shows samples’ low-dimensional embedding along with their canonical correlation with phenotype or MANOVA F-statistic (**Fig. 2C**). The ‘*treecor_degheatmap()*’ function can be used to show top differentially expressed genes of a given cell cluster (**Fig. 2E**). In order to summarize the final results, we also developed a versatile function *‘treecortreatplot()’* to display the multi-resolution cell type-phenotype association in a format we call ‘TreeCorTreat plot’ (**Fig. 2B,D,F**), which will be discussed next.

### TreeCorTreat plot for visualization

The TreeCorTreat plot is developed to provide a concise way to summarize the final analysis results at multiple resolutions. The plot consists of (1) a dendrogram showing the hierarchy of cell types and (2) information layers showing analysis results for each tree node. The ‘*treecortreatplot()*’ function provides a variety of display options for users to customize the plot and generate publication-quality figures. **Fig. 3** provides a few examples.

**Figure 3.**
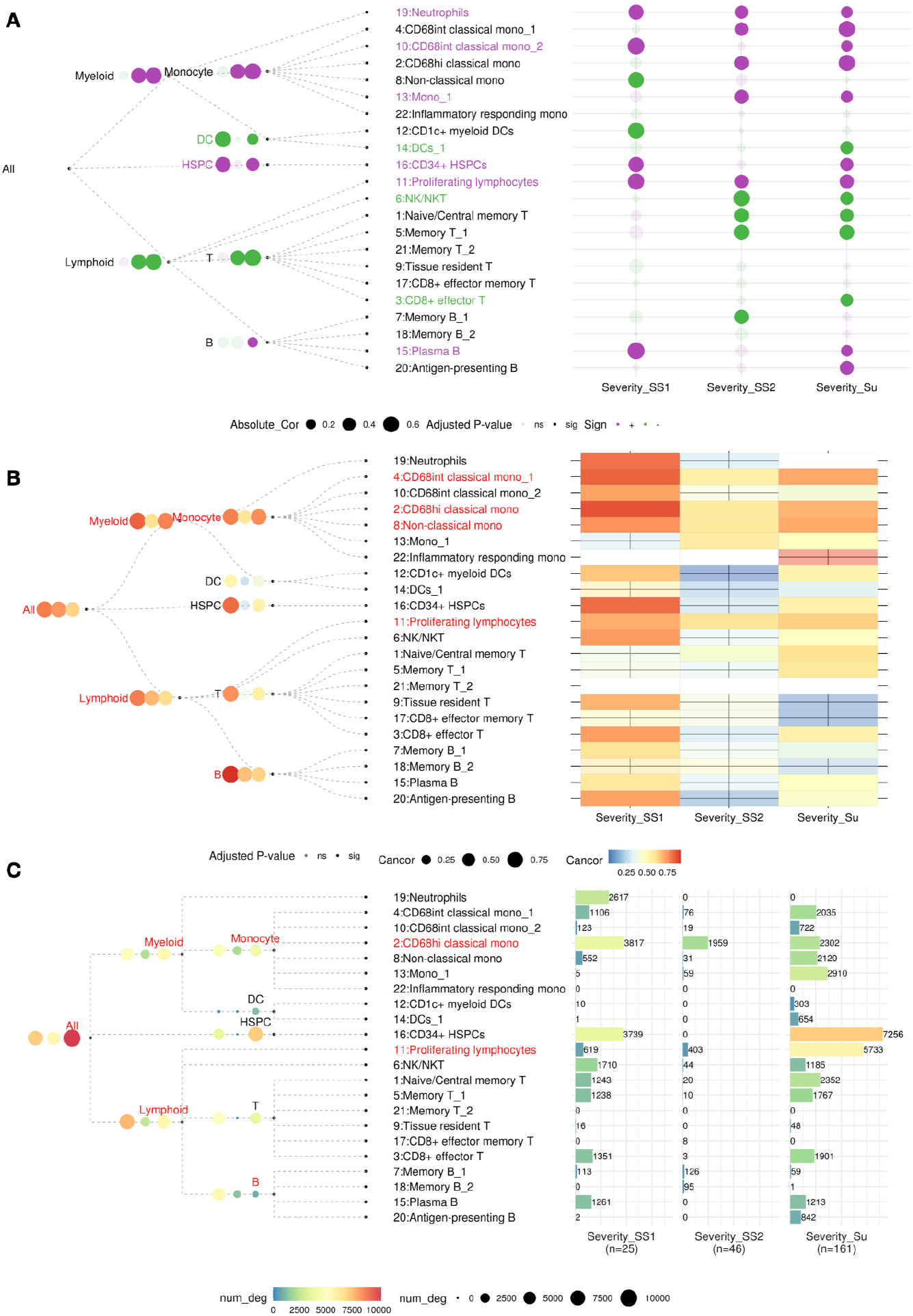
Assessment of study reproducibility when comparing mild to severe COVID-19 samples across three cohorts. (A) TreeCorTreat plot of cell type-sample phenotype association. For each tree node, three aligned circles correspond to results using SS1, SS2 and Su cohort respectively. In the balloon plot, circle size represents the magnitude of absolute value of Pearson correlation. Color indicates the sign of correlation, with green for negative (‘-’) and purple for positive correlation (‘+’). Transparency represents statistical significance after multiple testing corrections, with solid for statistically significant (‘sig’, adjusted p-value≤ 0.05) and transparent for non-significant (‘ns’, adjusted p-value>0.5). Colored labels highlight cell types with consistent patterns across three cohorts (purple for consistent positive correlation and green for consistent negative correlation). Here consistency is defined as showing statistically significant correlation in at least one cohort and having the same sign of correlation (i.e. positive or negative) across all three cohorts. For the root node ‘All’ and cell cluster ‘21:Memory T_2’, there is no correlation calculated because the cell type proportions are aggregated as 1 for all samples at root node and cluster 21 contains cells from only 1 sample in Su. (B) TreeCorTreat plot of global gene expression-sample phenotype association. In the left panel, both circle size and color represent the magnitude of canonical correlation. In the right panel (heatmap), color represents the magnitude of canonical correlation. Transparency represents statistical significance after multiple testing corrections, with transparent (see through) for non-significant (‘ns’, adjusted p-value>0.05) and solid for significant (‘sig’, adjusted p-value≤0.05). Red colored labels highlight cell types with consistent patterns across three cohorts. Here consistency is defined as showing statistically significant correlation in at least one cohort and having canonical correlation greater than 0.5 in all three cohorts. (C) TreeCorTreat plot of differential expression analysis. Color, circle size and bar height represent the number of DEGs. Red colored labels highlight cell types with reproducible patterns that have at least 100 differentially expressed genes identified in all three cohorts.

For displaying the tree dendrogram, the nodes can be connected using straight lines (**Fig.3A**), a quadratic bezier curve representation (**Fig.3B**), or a classical angle bend representation (**Fig.3C**). Users can choose line aesthetics such as line type (e.g. solid or dashed) and line color according to their preference.

For displaying the analysis results for each node, each TreeCorTreat plot consists of a non-leaf panel on the left and a leaf panel on the right. By default, balloon plots are used to display information for both leaf and non-leaf nodes. For the leaf panel, the rows in the balloon plot are aligned with leaf nodes in the dendrogram, and the plot can have multiple columns displaying different pieces of information. For the non-leaf panel, each internal tree node has a balloon plot next to it, showing information in a format similar to a leaf node.

As an example, in addition to the Su et al., we also analyzed the other two COVID-19 cohorts (SS1 and SS2). Since SS1 and SS2 do not have either healthy or moderate samples but all three cohorts have mild and severe samples (**Fig.2A**), we focused on comparing mild versus severe COVID-19 patients in this comparative analysis. **Fig.3A** shows the results of cell type proportion analysis in the three cohorts. At the finest cell type resolution, we observed consistent positive Pearson correlation with the disease severity in 6 cell types (i.e. purple cell clusters 10:CD68int classical mono_2, 11:Proliferating lymphocytes, 13:Mono_1, 15:Plasma B, 16:CD34+ HSPCs, 19:Neutrophils) and consistent negative correlation with the severity in 3 cell types (i.e. green cell clusters 3:CD8+ effector T, 6:NK/NKT, 14:DCs_1). Here ‘consistent’ is defined as showing statistically significant correlation in at least one cohort and having consistent sign of correlation (i.e. positive or negative) across all three cohorts. Such correlations likely are more replicable in other patient populations. By contrast, the correlation for the remaining cell clusters were either insignificant in all cohorts or inconsistent across the three cohorts.

In the balloon plots, users can use the size and color of the balloons to encode different information. In **Fig. 3A**, for instance, the balloon size reflects the magnitude of Pearson correlation, the color represents the sign (green: negative correlation; purple: positive correlation), and the transparency indicates the level of statistical significance (solid: adjusted p-value ≤ 0.05; transparent: adjusted p-value > 0.05). Users can customize these configurations according to their own preferences (e.g. use balloon size instead of transparency to show statistical significance, etc.). Users can also add texts to the balloon plot to show a value (e.g. correlation or adjusted p-value) for each balloon (**Additional file 5: Fig. S3**).

For the leaf panel, the balloon plot can also be replaced by other types of plot such as heatmap (**Fig. 3B**) and barplot (**Fig. 3C**). In **Fig. 3B**, the global gene expression analysis in the three COVID-19 cohorts shows that CD68+ classical monocytes, non-classical monocytes and proliferating lymphocytes showed consistent high canonical correlation (>0.5) with disease severity across all three cohorts in the mild versus severe disease comparison. In **Fig. 3C**, differential expression analysis shows that CD68 high classical monocytes and proliferating lymphocytes consistently showed >100 DEGs across all three cohorts in the mild versus severe comparison. In all these plot types, users can customize the text label, text color, layout (size/height/width), and display color configurations. For the non-leaf nodes, currently we do not provide options to replace balloon plots by other plot types due to the limited plotting space available within the tree dendrogram and the ensuing aesthetics challenges.

To the best of our knowledge, the ‘*treecortreatplot()*’ function is the first tool that allows users to programmatically generate TreeCorTreat plots in one command. Its ability to automatically align the dendrogram with information plots (i.e. balloon plot, heatmap, etc.) and its support for a wide range of customizable configurations makes it flexible and adaptable to displaying information in many different applications.

### TreeCorTreat supports versatile multi-resolution analysis

To analyze cell type-phenotype association at multiple resolutions, fine-grained cell clusters (children) are progressively merged into bigger clusters (parents). When a parent node is created by merging two or more child nodes, the information from the children can be combined in multiple ways with different implications. The two basic methods to combine child nodes are ‘aggregation’ and ‘concatenation’. Consider the toy example in **Figs. 4A** and **4B** where two child nodes are combined into a parent node. The aggregation method pools all cells from the child nodes into a single pseudobulk sample which is used to represent the parent node (**Fig. 4A**). In the concatenation method, one first pools cells within each child node to form a pseudobulk sample for that child node. Each child node is then represented using a feature or a feature vector extracted from its pseudobulk sample (e.g., highly variable genes). Next, the parent node is represented by concatenating the feature vectors from all child nodes into a longer vector (**Fig. 4B**). In the toy example, column vectors ***v***_1_ and ***v***_2_ are the gene expression vectors for the two children respectively, and suppose *p*_1_ and *p*_2_ are the cell type proportion of the two child nodes in a sample, then the gene expression vector for the parent node will be (*p*_1_***v***_1_ + *p*_2_***v***_2_)/(*p*_1_ + *p*_2_) up to a proportionality constant in the aggregation method, and it will be [***v***_1_’, ***v***_2_’]’ in the concatenation method.

**Figure 4.**
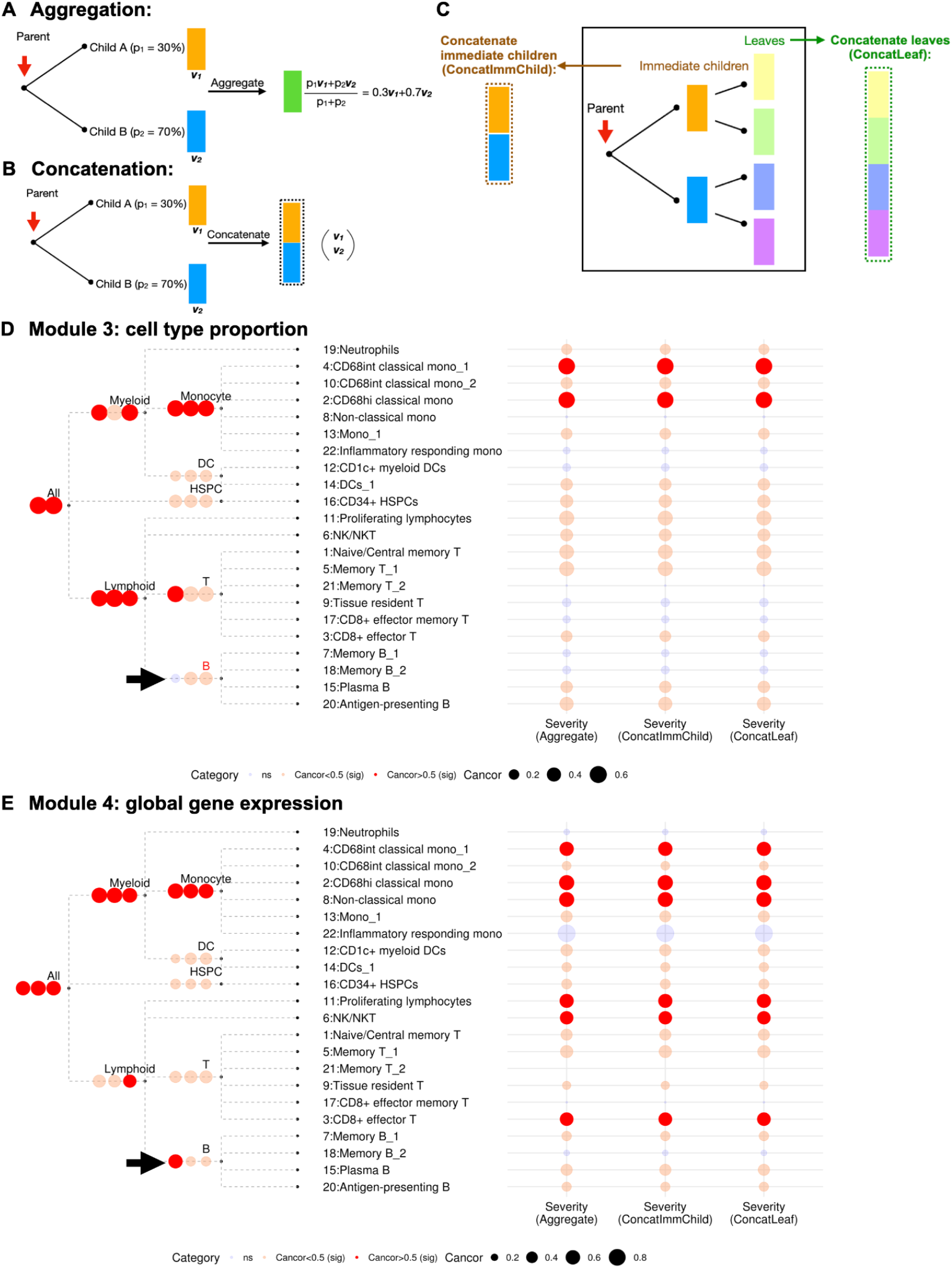
Comparison of three feature summary approaches, using a single COVID-19 dataset (Su et al.) as a demonstration. (A) Cartoon for aggregating features from child nodes to a parent node. (B) Cartoon for concatenating child-specific features at a parent node. (C) Cartoon of two concatenation subtypes, with concatenating immediate children on the left (‘ConcatImmChild’ labelled in brown) and concatenating leaf nodes on the right (‘ConcatLeaf’ labelled in green). (D) Comparison of three feature summary approaches in cell type proportion-disease severity association. For each tree node, three aligned circles correspond to results using ‘Aggregate’, ‘ConcatImmChild’ and ‘ConcatLeaf’ approaches respectively. Circle size represents the magnitude of canonical correlation and color indicates three categories defined by multiple testing adjusted p-value and effect size (blue for non-significant (‘ns’), light red for significant (‘sig’) with canonical correlation<0.5, and solid red for significant (‘sig’) with canonical correlation>0.5). Here canonical correlation instead of Pearson correlation is shown for the ‘Aggregate’ approach for convenience of comparison with the other two methods. Red colored label with an arrow highlights cell type with a change in statistical significance using three merging approaches. For the root node, there is no correlation for the ‘Aggregate’ approach because the cell type proportions are aggregated as 1 for all samples. (E) Comparison of three feature summary approaches in global gene expression-disease severity association, with the same color palette as shown in Fig.4D.

To support users’ diverse needs, we further provide two concatenation options. In the ‘concatenate leaves’ option, feature vectors of all terminal leaf nodes derived from a node are concatenated into a long vector to serve as the feature vector of the node (**Fig.4C**). In the ‘concatenate immediate children’ option, a node’s immediate children are first identified. The feature vector of each immediate child is obtained using the aggregation approach. Then the feature vector of the node in question is obtained by concatenating the feature vectors of its immediate children (**Fig.4C**).

All three ways to combine child nodes (aggregate, concatenate leaves, and concatenate immediate children) are provided for both the cell type proportion analysis and global gene expression analysis. For the cell type proportion analysis, the concatenation may produce a feature vector (i.e., proportions of multiple child cell types) rather than a univariate feature (i.e. the proportion of one single cell type). When the feature is a multivariate vector, the Pearson or Spearman’s correlation between the feature and phenotype will be replaced by the canonical correlation obtained using CCA. For differential gene expression analysis, the feature for each node is its pseudobulk gene expression profile obtained using the aggregation approach. Here concatenation is redundant and unnecessary since differential genes for the leaf nodes and immediate children are all already reported in the aggregation approach.

When analyzing how the parent node is associated with the phenotype, the concatenation method only considers the combined effect of child nodes’ features (e.g., gene expression of each child node) without considering their relative cell type proportions (i.e. how different child cell types are mixed together). By contrast, the aggregation method considers how the children are mixed together as well. If the parent node is associated with the phenotype using the ‘aggregation’ approach, the association could be either due to the mixing proportions of child nodes being associated with the phenotype, or due to the children’s own features such as childrens’ gene expression profiles being associated with the phenotype, or due to both. On the other hand, if the association is found using the ‘concatenation’ approach, the association observed at the parent node is due to the association between the phenotype and children’s features such as childrens’ gene expression profiles, and it is not due to the association with the mixing proportions of child cell types in different samples.

As a conceptual example, consider four leaf nodes with the following cell type proportion in a sample: Monocytes (0.2), Granulocytes (0.05), T cell (0.2) and B cell (0.1). Suppose T cell and B cell are merged into an intermediate node ‘Lymphocytes’, and Lymphocytes, Monocytes and Granulocytes are further merged into a node ‘White Blood Cell’ (**Additional file 5: Fig. S4**). For the node ‘White Blood Cell’, the feature defined by ‘aggregation’ will be 0.55, the feature defined by ‘concatenate leaves’ will be a vector [0.2, 0.05, 0.2, 0.1], and the feature defined by ‘concatenate immediate children’ will be a vector [0.2, 0.05, 0.3]. In practice, the way to define features for the intermediate and root nodes depends on the analysis needs. For example, If one is interested in whether the white blood cell proportion in a sample is associated with the sample’s phenotype, then one should use ‘aggregation’. On the other hand, if one is interested in whether the white blood cells are associated with phenotype via the mixing proportions of lymphocytes, monocytes and granulocytes, then one should use ‘concatenate immediate children’. Finally, if one is further interested in whether the white blood cells are associated with phenotype via mixing proportions of T cells, B cells, monocytes and granulocytes, then one should use ‘concatenate leaves’.

As a real data example, we reanalyzed the association between COVID-19 disease severity and cell type proportion in the Su et al. using all three merging methods (**Fig.4D**). For most cell types, the phenotype-cell type proportion association remained similar across different merging methods. However, for B cells, using the ‘aggregation’ approach resulted in insignificant association, whereas using ‘concatenate leaves’ and ‘concatenate immediate children’ resulted in statistically significant association between B cells and disease severity. This suggests that, while the proportion of the aggregated B cell population is not significantly correlated with severity, the relative abundance of B cell subtypes within the B cell population is associated with disease severity.

Similarly we also rerun the global gene expression analysis using all three merging methods (**Fig.4E**). Overall, the phenotype-global expression association remained similar across different merging methods for most cell types. For B cells, using ‘concatenate leaves’ and ‘concatenate immediate children’ both resulted in statistically significant association between B cells and severity. However, using the ‘aggregation’ approach resulted in stronger correlation with disease severity, suggesting that the gene expression profiles of all B cell subtypes combined with the cell subtype mixing proportions correlate with disease severity better than the gene expression profiles of B cell subtypes alone. This is also consistent with **Fig. 4D** which shows that the mixing proportions of B cell subtypes within the B cell population is associated with disease severity.

### Analysis of multivariate outcomes

When there are multiple phenotypic traits, one can either analyze each trait separately as a univariate phenotype or analyze them jointly as a multivariate phenotype. For analyzing association between a multivariate phenotype and cell type proportion or global gene expression, by default CCA is used to compute the canonical correlation between the phenotype and cell type features (i.e. cell type proportion or gene expression). Statistical significance is determined using permutation test p-values after adjusting for multiple testing.

Optionally, one can also transform a multivariate phenotype into a univariate phenotype (e.g., using principal component analysis, or a weighted linear combination of traits, or user-defined ways to categorize samples into a binary or categorical variable by jointly considering multivariate traits) and then run the TreeCorTree analyses developed for univariate phenotype.

For analyzing differential gene expression, a multivariate phenotype is first transformed into a univariate phenotype using either the principal component analysis, or a weighted linear combination of traits, or user-defined ways to categorize samples. The transformed univariate phenotype, which combines information from multiple traits, is then used to run limma differential expression analysis. **Table 1** summarizes the TreeCorTreat analysis procedures for different scenarios.

**Table 1.** Summary table for TreeCorTreat analyses.

| Module | Phenotype | Methods to summarize features | Methods to analyze association |  |
| --- | --- | --- | --- | --- |
|  |  |  | Leaf node | Non-leaf node |
| Cell type proportion | Univariate | Aggregate | PCC or SCC or regression | PCC or SCC or regression |
|  |  | Concatenate leaf | CCA or regression | PCA+CCA or PCA+regression |
|  |  | Concatenate immediate children | CCA or regression | PCA+CCA or PCA+regression |
|  | Multivariate (separate) | Aggregate | PCC or SCC or regression | PCC or SCC or regression |
|  |  | Concatenate leaf | CCA or regression | PCA+CCA or PCA+regression |
|  |  | Concatenate immediate children | CCA or regression | PCA+CCA or PCA+regression |
|  | Multivariate (joint) | Aggregate | CCA or regression | CCA or regression |
|  |  | Concatenate leaf | CCA or regression | PCA+CCA or PCA+regression |
|  |  | Concatenate immediate children | CCA or regression | PCA+CCA or PCA+regression |
| Global gene expression | Univariate | Aggregate | HVG+PCA+CCA or HVG+PCA+regression | HVG+PCA+CCA or HVG+PCA+regression |
|  |  | Concatenate leaf | HVG+PCA+CCA or HVG+PCA+regression | HVG+PCA+CCA or HVG+PCA+regression |
|  |  | Concatenate immediate children | HVG+PCA+CCA or HVG+PCA+regression | HVG+PCA+CCA or HVG+PCA+regression |
|  | Multivariate (separate) | Aggregate | HVG+PCA+CCA or HVG+PCA+regression | HVG+PCA+CCA or HVG+PCA+regression |
|  |  | Concatenate leaf | HVG+PCA+CCA or HVG+PCA+regression | HVG+PCA+CCA or HVG+PCA+regression |
|  |  | Concatenate immediate children | HVG+PCA+CCA or HVG+PCA+regression | HVG+PCA+CCA or HVG+PCA+regression |
|  | Multivariate (joint) | Aggregate | HVG+PCA+CCA or HVG+PCA+regression | HVG+PCA+CCA or HVG+PCA+regression |
|  |  | Concatenate leaf | HVG+PCA+CCA or HVG+PCA+regression | HVG+PCA+CCA or HVG+PCA+regression |
|  |  | Concatenate immediate children | HVG+PCA+CCA or HVG+PCA+regression | HVG+PCA+CCA or HVG+PCA+regression |
| Differential expression | Univariate | / | limma | limma |
|  | Multivariate (separate) | / | limma | limma |
|  | Multivariate (joint) | / | limma | limma |

Note: PCC - Pearson correlation coefficient; SCC - Spearman correlation coefficient; HVG - selecting highly variable genes; PCA - principal component analysis; CCA - canonical correlation analysis.

To demonstrate, consider adding age to the COVID-19 analysis using the Su et al. COVID-19 disease severity has been reported to be associated with age [40,61,62]. We first treat disease severity and age as two separate phenotypes. We asked whether they are associated with similar cell type features. In the global gene expression analysis, we found a total of 17 out of 30 cell types at different resolutions that are associated with both the severity and age (**Fig. 5A,** cell types with red text labels).

**Figure 5.**
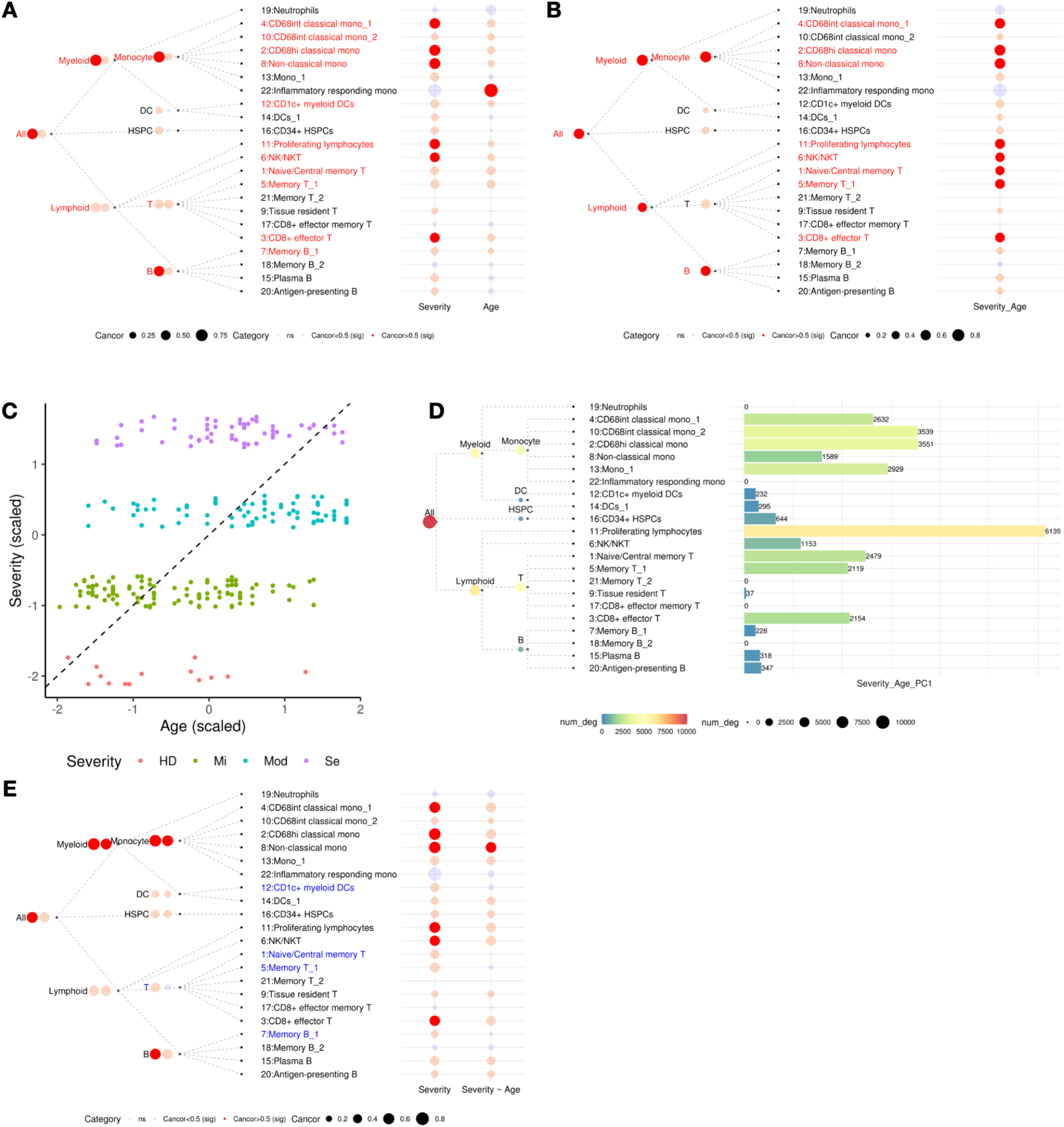
Analysis on multivariate phenotypes in Su COVID-19 dataset. (A) TreeCorTreat plot of association between global gene expression and disease severity or age separately. Circle size represents the magnitude of canonical correlation and color indicates three categories defined by multiple testing adjusted p-value and effect size (blue for non-significant (‘ns’), light red for significant (‘sig’) with canonical correlation<0.5, and solid red for significant (‘sig’) with canonical correlation>0.5). Red colored labels highlight cell types with significant associations with both severity and age. (B) TreeCorTreat plot of association between global gene expression and joint (disease severity, age) phenotype. Red colored labels highlight statistically significant cell types with canonical correlation>0.5. (C) Scatter plot of scaled severity against scaled age with dashed line representing first principal component (PC1) axis. (D) Differential expression analysis of PC1 of age and severity as the phenotype (dashed line in (C)). Color, circle size and bar height represent the number of differentially expressed genes. (E) Comparison of global gene expression-severity association before and after adjusting for age as a covariate, with the same color palette as shown in (A)-(B). Blue colored labels highlight cell types with changes in multiple testing adjusted p-value from significant (light red or solid red) to non-significant (blue) after adjusting for age.

We then analyzed severity and age jointly as a two-dimensional phenotype and asked which cell types are associated with this joint phenotype. We found that 25 out of 30 cell types are significantly associated with the joint phenotype (**Fig. 5B,** dark and light red balloons). Among them, 13 cell types had canonical correlation larger than 0.5 (**Fig. 5B,** dark red balloons). We also extracted the first principal component (PC1) of severity and age (**Fig. 5C**) and identified differentially expressed genes associated with PC1 (**Fig. 5D**). In both analyses, CD68+ classical monocytes, non-classical monocytes, proliferating lymphocytes, NK/NKT cells, naive and memory T cells, and CD8+ effector T cells showed the strongest gene expression changes associated with the joint phenotype (**Fig. 5B,D**).

### Adjusting for covariates

TreeCorTreat can handle covariates in the analysis. This allows users to adjust for potential confounders or unwanted technical variation. For example, instead of viewing age as a phenotype, one can also view it as a covariate and ask which cell type features are associated with disease severity after accounting for age. **Fig. 5E** shows such an analysis on global gene expression using the same Su dataset as in **Fig. 5A-D**. After adjusting for age, the association between the COVID-19 severity and four leaf cell types (CD1c+ myeloid DCs, naive/central memory T cells, memory T_1, memory B_1) and one intermediate cell type (T cell) became insignificant. By contrast, the other severity-associated cell types, such as CD68+ classical monocytes, non-classical monocytes, proliferating lymphocytes, NK/NKT cells, and CD8+ effector T cells, etc., are associated with severity both before and after adjusting for age (**Fig. 5E**).

As another example, we performed global gene expression analysis using all samples from the three COVID-19 cohorts together (coding severity as HD = 1, Mi = 2, Mod = 3, Se = 4). To do so, one has to first remove batch effects. Taking cluster 4 (CD68 intermediate classical monocytes) as an example, **Fig. 6A** shows the low-dimensional embedding of samples in the first two PCs of the gene expression of this cell type. Samples are separated based on cohort, indicating clear batch effects. In our TreeCorTreat analysis, we fitted linear regression and added cohort identifiers (study ID) as potential confounders. **Fig. 6B** shows the samples’ embedding after regressing out the batch effects by including study ID as covariates in regression. The batch effects are largely removed and samples from different cohorts are mixed well. The residuals of the regression were then used to perform highly variable gene selection, PCA and CCA. After adjusting for study ID (i.e. batch), 2 intermediate nodes (i.e. lymphoid and T cells) and 9 leaf cell types (neutrophils, two CD68 intermediate classical monocyte clusters, CD1c+ myeloid DCs, NK/NKT, native/central memory T, memory T_1, CD8+ effector memory T, and CD8+ effector T cells) changed their statistical significance from nonsignificant to significant (adjusted p-value≤0.05; **Fig. 6C**). The CD68 intermediate classical monocyte example again illustrates how this change in significance occurred: the batch effects dominated the gene expression variability before adjustment and obscured its association with severity (**Fig. 6A**), whereas the association with severity became clear after removing the batch effects (**Fig. 6B**).

**Figure 6.**
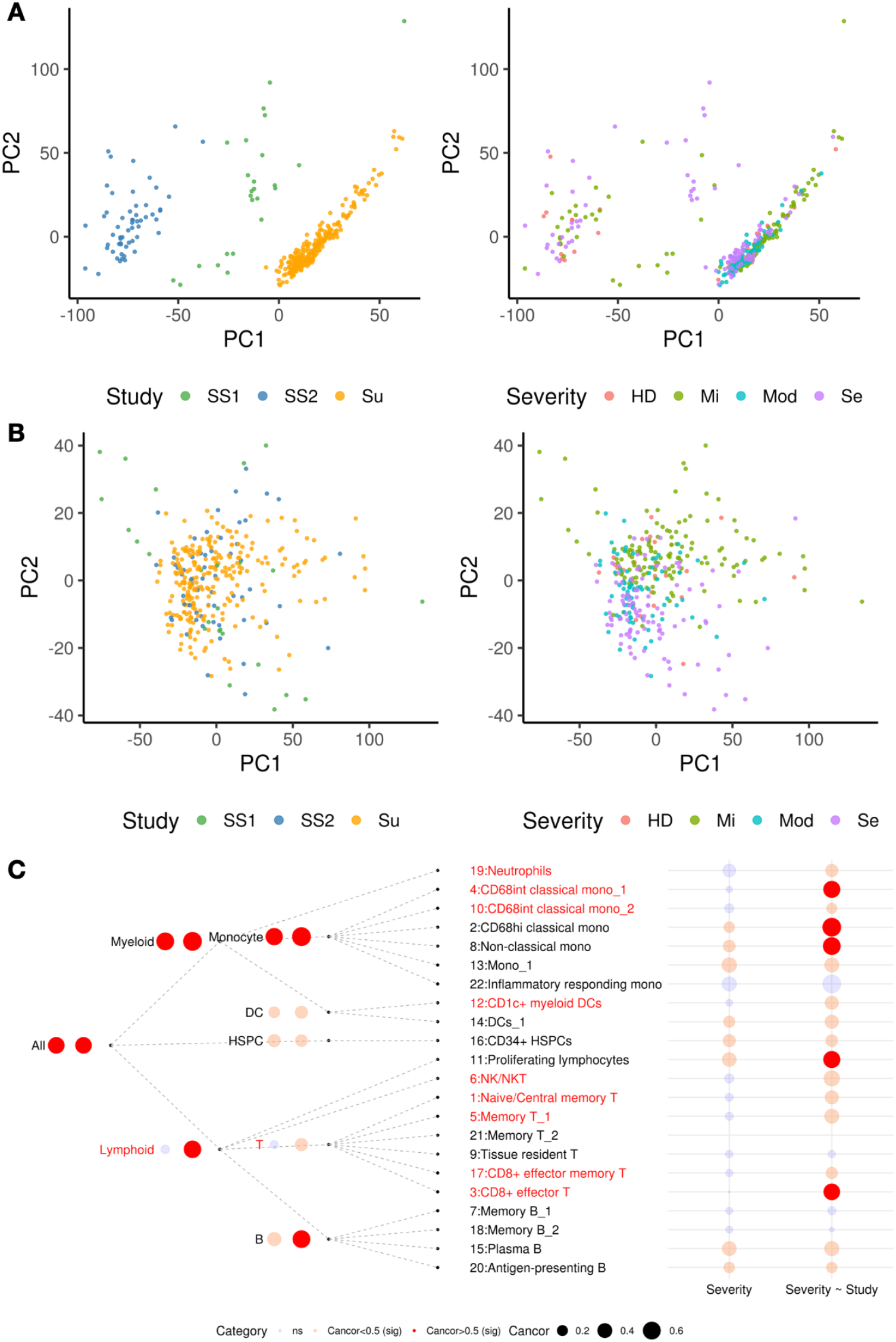
Assessment of batch effect in the combined analysis of SS1, SS2 and Su COVID-19 cohorts. (A) PCA plot of sample-level pseudobulk gene expression profile for cell cluster ‘4: CD68 intermediate classical monocytes’ with color coding for cohort ID (left) and disease severity (right). (B) PCA plot of batch corrected sample-level pseudobulk expression profile for cell cluster ‘4: CD68 intermediate classical monocytes’ with color coding for cohort ID (left) and disease severity (right). (C) Comparison of association between global gene expression and disease severity before (1st column) and after batch adjustment (2nd column). Here three cohorts are combined and treated as one dataset. Circle size represents the magnitude of canonical correlation and color indicates three categories defined by multiple testing adjusted p-value and effect size (blue for non-significant (‘ns’), light red for significant (‘sig’) with canonical correlation<0.5, and solid red for significant (‘sig’) with canonical correlation>0.5). Red colored labels highlight cell types that changed from non-significant to significant after adjusting for cohort ID.

### Computational time and memory usage

**Fig. 7** and **Additional file 2: Table S2** summarize the run time and memory (RAM) usage of each module in TreeCorTreat using the COVID data as an example. The analyses were run on either the Su dataset alone (253 samples, 500,780 cells) or all three cohorts jointly (335 samples, 659,375 cells). The *treecor_harmony* module (module 1), which consists of Harmony integration, louvain clustering and cell type specific differential gene marker identification, is the most time consuming and RAM intensive module (**Fig. 7A**). It took ~3.5 days and ~80GB RAM for analyzing the Su et al dataset and 5.5 days and ~120GB RAM for analyzing the three cohorts jointly using a single CPU core (2.5 GHz). Since this module calls ‘*RunHarmony*’, ‘*FindClusters*’ and ‘*FindAllMarkers*’ functions in Seurat, its speed and memory usage mainly depend on these external functions and packages.

**Figure 7.**
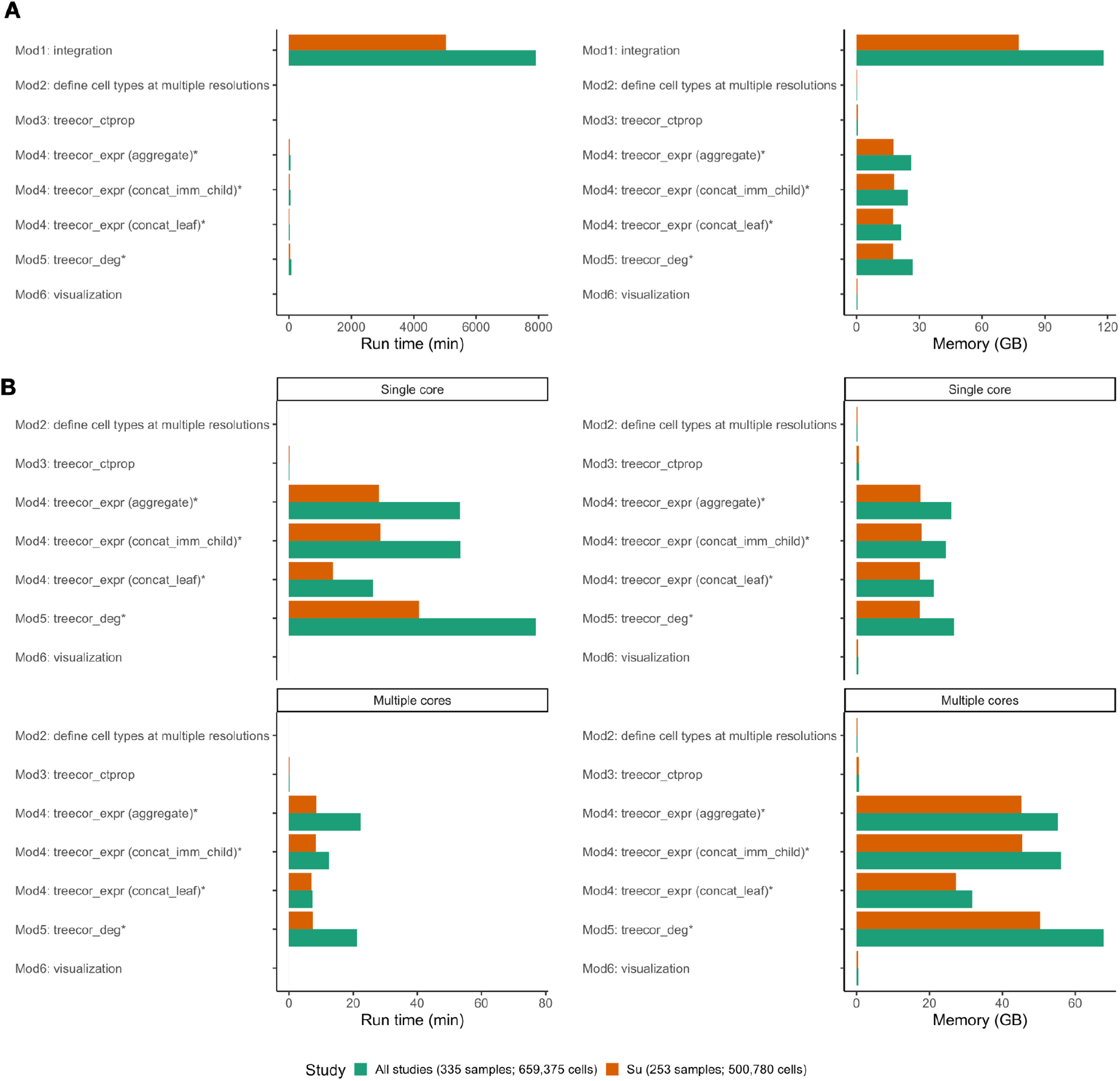
Computational time and memory usage for each TreeCorTreat module. (A) A summary of computational time in minutes (left) and memory usage in gigabytes (GB) (right). (B) Excluding data integration (module1), summary of computational time in minutes (left) and memory usage in GB (right) of remaining five modules, using either single-core (upper panel) or multi-core implementations (lower panel). Orange represents the performance of running a single study (Su, 253 samples and 500,780 cells) and green represents the performance on all studies combined (335 samples and 659,375 cells).

Excluding the *treecor_harmony* module, running all the other modules in a single CPU core as above took a total of 111 minutes and a maximum of 17.91 GB RAM for the Su dataset, and it took a total of 210 minutes and maximum of 26.71 GB RAM for analyzing all three cohorts jointly. For these modules, the computation can be accelerated by using multiple cores, although with increased total RAM usage. On 24 cores, the computation for the Su dataset took a total of 32 minutes and a combined maximum of 50.39 GB RAM, and it took a total of 63 minutes and a combined maximum of 67.80 GB RAM for jointly analyzing all three cohorts (**Fig. 7B**).

### TreeCorTreat analysis of scRNA-seq data from non-small cell lung cancer (NSCLC)

As a second real data example, we applied TreeCorTreat to a recently published scRNA-seq dataset from neoadjuvant anti-PD-1 (nivolumab) treated non-small cell lung cancers (NSCLC)[43]. This dataset contains 31 samples from 16 treated patients, 9 of whom were defined as responders based on clinical assessment. Among 31 samples, there are 15 tumour-infiltrating lymphocyte (TIL) samples and 12 adjacent normal lungs, 11 of which are paired TIL and normal samples. We obtained the integrated data from the original publication, where the integration was done using Seurat and 15 distinct T cell clusters were defined and annotated based on top differentially expressed genes or known immune markers. From the integrated data, we obtained a total of 469,418 T cells from the 15 TIL and 12 normal lung samples and used them for the TreeCorTreat analysis (**Fig.8A**). The 15 cell clusters include a MAIT cell cluster, 7 CD8 T cell clusters (i.e. CD8-TRM-LINC02446, CD8-TRM-TYROBP, CD8-Effector-TYROBP, CD8-Effector-GZMK, CD8-Effector-CCL4L2, CD8-Proliferating and MHCII) and 7 CD4 T cell clusters (i.e. CD4-Naive/Stem like-SELL, CD4-Th-PLIN2, CD4-Th-MT1E, CD4-Th-IL7R, CD4-Tfh-CXCL13-1, CD4-Tfh-CXCL13-2 and CD4-Treg).

**Figure 8.**
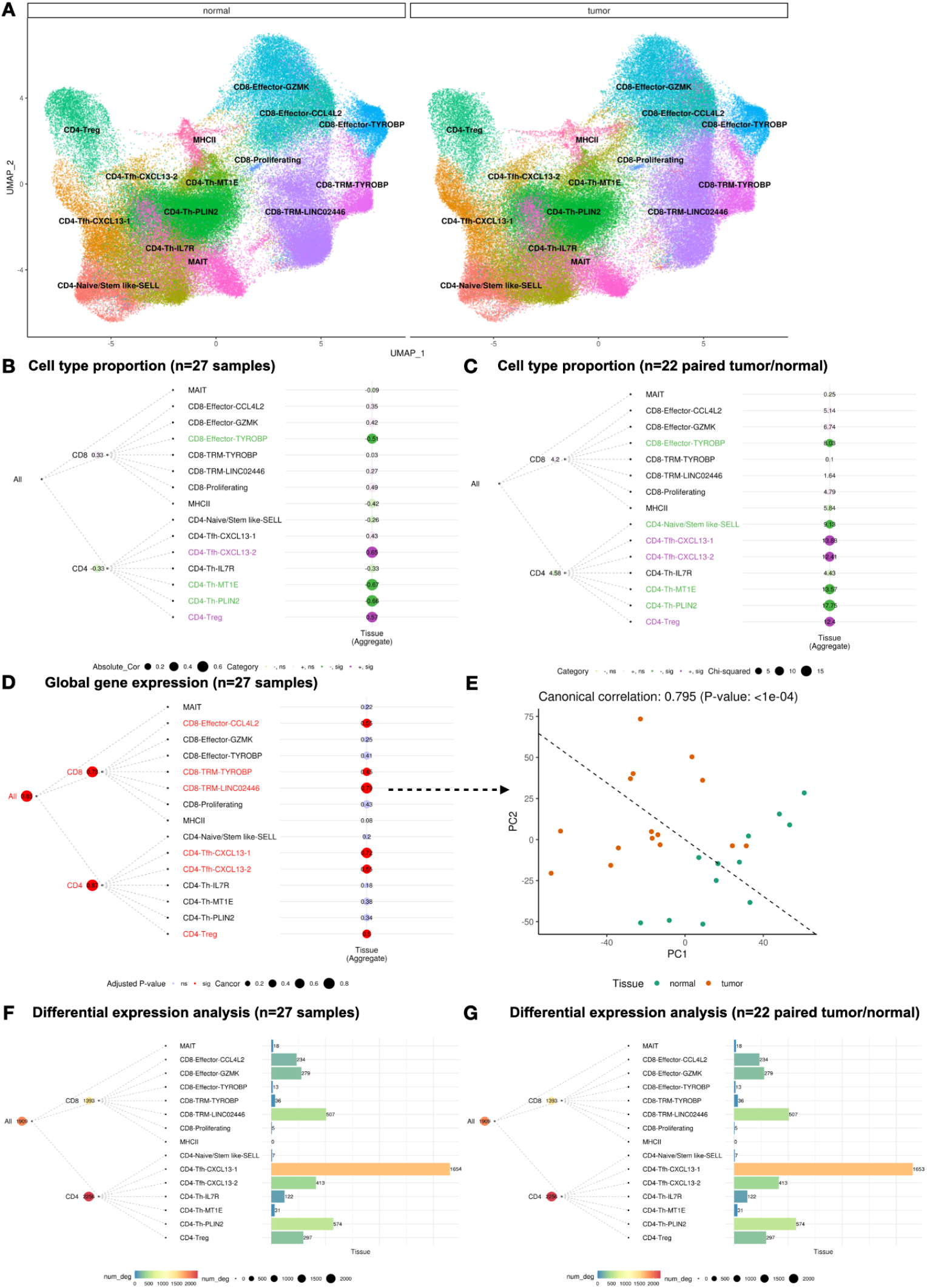
TreeCorTreat analyses of the non small cell lung cancer (NSCLC) dataset. (A) UMAP of 15 cell clusters stratified by tissue type (left: normal lung; right: tumor). (B) TreeCorTreat plot of cell type-tissue association with default setting (aggregate) using unpaired 27 samples (15 tumors and 12 normals). In the balloon plot, circle size represents the magnitude of absolute value of Pearson correlation. Colors indicate four categories defined by correlation sign (‘+’ or ‘-’) and statistical significance (‘sig’ or ‘ns’), with green for negative correlation, purple for positive correlation, dark for significant, and transparent for nonsignificant. Colored labels highlight cell types with significant correlation after multiple testing correction. (C) TreeCorTreat plot of cell type-tissue association with default setting (aggregate) using 22 samples (11 matching pairs of tumor and normal samples), with the same color palette in (B). Circle size corresponds to Chi-squared likelihood ratio statistic, comparing a full and a reduced model with and without including tissue as an independent variable. (D) TreeCorTreat plot of global gene expression-tissue association with default setting (aggregate) using 27 unpaired samples. Circle size corresponds to canonical correlation and color indicates statistical significance: blue for non-significant (‘ns’) and red for significant (‘sig’, adjusted p-value≤0.05). (E) PCA plot of sample-level pseudobulk in ‘CD8-TRM-LINC02446’ cell type for individual samples with color annotating tissue compartments (green for normal, orange for tumor). The dashed line represents the optimal axis inferred from CCA. For (B)-(E), p-value was determined by permuting sample labels 10,000 times and the BY procedure was used for multiple testing adjustment. (F) Differential expression analysis of tissue type using 27 unpaired samples. (G) Differential expression analysis of tissue type using 22 samples (11 matching pairs of tumor and normal samples).

We asked how T cells in normal lung and tumor are different (let normal = 0 and TIL = 1). Since the patients were collected from two different cancer centers, we regressed out potential batch effects by treating cancer center ID as a covariate. The default cell type proportion analysis (‘Aggregate’, Pearson correlation) shows that cell types most correlated with tissue compartment include CD4-Tfh-CXCL13-2 (a CD4 T follicular helper cell subpopulation with high *CXCL13* expression) and CD4-Treg (CD4 regulatory T cells), which have higher proportion in TIL than in normal lung, and CD8-Effector-TYROBP (CD8 effector T cells with high *TYROBP* expression), CD4-Th-MT1E (CD4 T helper cell with high *MT1E* expression) and CD4-Th-PLIN2 (CD4 T helper cell with high *PLIN2* expression), which have lower proportion in TIL than in normal lung (**Fig.8B**). Since the default analysis does not consider the sample pairing, we further focused our analysis on the 11 patients with matched TIL and normal samples. We run the regression analysis for each tree node that considers the matching by additionally introducing patient indicators as covariates, and we asked whether the result in the unpaired Pearson correlation analysis (27 samples in total) is robust and remains true in this paired analysis (22 samples in total). **Fig. 8C** shows that all aforementioned patterns observed in unpaired analysis hold true in the paired analysis. The paired analysis additionally identified CD4 naive/stem-like cells with high *SELL* expression (CD4-Naive/Stem like-SELL) as a cluster with significantly lower proportion in TIL than in normal lung. We also performed analyses using the ‘Concatenate leaves’ and ‘Concatenate immediate children’ options and found largely similar results (**Additional file 5: Fig. S5**).

In terms of gene expression, three parent cell types (i.e All, CD4, CD8) and six leaf cell types (i.e. CD8-Effector-CC4L2, CD8-TRM-TYROBP, CD8-TRM-LINC02446, CD4-Tfh-CXCL13-1, CD4-Tfh-CXCL13-2, CD4-Treg) showed significant (adjusted p-value≤0.05) and strong canonical correlation (>0.5) with the tissue compartment in the global gene expression analysis (**Fig.8D**; ‘Aggregate’). T cells from tumor and normal lung samples can indeed be well separated based on gene expression in these cell types, with an example shown for CD8-TRM-LINC02446 (**Fig.8E**). Similar cell types were identified using the ‘Concatenate leaves’ or ‘Concatenate immediate children’ options (**Additional file 5: Fig. S6**). The global expression analysis provides a way to visualize samples in the low-dimensional PC space, but it does not generate the list of differential genes or consider the sample pairing. Therefore, we proceeded with the differential expression analysis. When the differential expression analysis was run by treating 15 TIL and 12 normal lung samples as two unpaired groups, CD8-Effector-CC4L2, CD8-Effector-GZMK, CD8-TRM-LINC02446, CD4-Tfh-CXCL13-1, CD4-Tfh-CXCL13-2, CD4-Th-PLIN2 and CD4-Treg cells showed the largest number of DE genes (**Fig.8F**). When the analysis was run on the 11 matched pairs of TIL and normal lung samples using a paired test, the results remained largely similar **(Fig.8G)**.

### Comparison with existing tools

As illustrated above, analyzing a rich multi-sample scRNA-seq dataset to understand how different cell types are linked to samples’ phenotype is a complex task. It often requires one to explore different feature types at multiple cell type resolutions, examine impacts of covariates, rule out association due to potential confounders, investigate links among different traits, compare results from different analysis protocols to assess robustness of the results, and evaluate consistency across datasets. This means that one often needs to run and compare dozens to hundreds of different analyses on his/her huge dataset in order to identify the most important and reliable cell type-phenotype associations. For example, in our COVID analysis, certain associations are consistently found in multiple datasets (e.g. the association between severity and cell proportions of monocytes and T cells, and gene expression of CD68+ classical monocytes and proliferating lymphocytes; **Fig. 3**) and across different analyses (e.g. the association between severity and gene expression of CD68+ classical monocytes and proliferating lymphocytes with or without adjusting for age in the Su dataset in **Fig. 5E** and before or after correcting for batch effects in the joint analysis of all three COVID cohorts in **Fig. 6C**). Once these robust associations are identified, they can help one to focus the available resources on studying the most promising cell and feature types in the in-depth downstream computational and experimental studies. By contrast, some associations depend on the analysis protocol (e.g. the association between COVID-19 severity and CD1c+ myeloid DCs, naive/central memory T cells, memory T_1, memory B_1 depends on whether one adjusts for age; **Fig. 5E**), and correcting for batch effects or not will also affect the results when jointly analyzing multiple cohorts; **Fig. 6C**). In all these explorations, being able to visualize the results is important for comparing different analyses and digesting the large amounts of information.

This exploration cannot be efficiently carried out using existing bulk DE analysis tools such as limma [53], DESeq [54,55] and edgeR [56]. These tools do not automatically identify cell subpopulations in each sample and do not run the analysis for each cell subpopulation and at multiple cell type resolution levels as in **Figs. 2–4**. Similarly, commonly used single-cell DE analysis methods such as Seurat [17], MAST[49], SCDE[50], scDD[51] and DEsingle[52] also do not perform analysis at multiple resolutions and do not support different ways to define and merge features along the cell type hierarchy as in **Figs. 2–4**. Moreover, they are mainly designed for comparing two cell populations or two samples and cannot handle complex multi-sample comparison problems where samples’ phenotype can be univariate or multivariate and there might be covariates that need to be adjusted for, as in **Figs. 5–6.** None of the above tools provides a visualization tool like the TreeCorTreat plot to help data analysts conveniently compile and visually compare the results which is important for extracting big pictures from the large amounts of analysis results. While it is not impossible to handle some of the analysis tasks manually (e.g. run limma for each cell cluster manually to detect DE genes, or draw the TreeCorTree plot manually), this manual approach is not scalable to running a large number of exploratory analyses on a large number of cell clusters at many different resolution levels. Alternatively, data analysts could develop their own codes to do the analyses. However, even for an experienced data analyst and programmer, the coding, debugging, implementing different analysis options (e.g. different ways to merge child nodes, different feature types, etc.), and finding optimal ways to visualize data will take many days of time and substantial amounts of efforts, and it is therefore highly inefficient if each data analyst has to develop his/her own code from scratch. By contrast, TreeCorTreat provided a wide variety of analysis functions, whose R commands along with usage examples are summarized in **Additional file 3: Table S3**, to support a systematic exploration of phenotype-cell type associations. It also provides TreeCorTreat plot as a new visualization tool. As multiple sample scRNA-seq data become increasingly common and identifying phenotype-cell type association is one of the most common analyses for such data, TreeCorTreat will save vast amounts of data analysts’ time which can be redirected to tackling other important problems towards understanding biology. As such, TreeCorTreat will help accelerate scientific discovery using multi-sample scRNA-seq data.

## Discussion

As this article demonstrates, TreeCorTreat provides a software tool that makes the exploration and visualization of cell type-phenotype association in multi-sample scRNA-seq datasets easier and more accessible to data analysts. The cell types and feature types most relevant to the phenotype identified by TreeCorTreat can provide targets for in-depth downstream analyses. For example, once a cell type is found to be associated with phenotype in terms of gene expression, one can take a closer look at the differentially expressed genes for that cell type to further identify promising gene candidates or enriched pathways using a variety of existing tools such as gene set enrichment analysis [63] or gene ontology analysis [64–68] to develop mechanistic insights into the phenotype-cell type association.

Currently, TreeCorTreat is primarily designed for applications where a sample consists of multiple cell subpopulations and one wants to analyze phenotype association for each cell subpopulation. The current implementation of TreeCorTreat does not consider applications where cells in a sample form a continuous trajectory and one wants to analyze the association between sample phenotype and gene’s dynamic expression program along the trajectory. For that purpose, recent methods such as condiments[69] and Lamian[70] may be used. Lamian, in particular, can analyze multiple samples using a general regression framework. Condiments, on the other hand, can compare two different conditions by assuming that each condition has one sample.

In the future, TreeCorTreat may be extended in multiple directions. First, we will continue to add new methods for analyzing the phenotype-cell type association to the software, such as new scRNA-seq differential gene expression analysis methods as they emerge. Second, while TreeCorTreat currently only supports scRNA-seq data, its general analysis and visualization framework may be used to handle other single-cell genomic data types as well. For example, one may develop a TreeCorTreat pipeline for single-cell ATAC-seq or single-cell multiomics data after tailoring the method details to account for data type specific characteristics. These extensions will be an important topic for future investigation.

## Conclusions

In summary, TreatCorTreat provides a solution to analyzing and visualizing multi-resolution phenotype-cell type association in multi-sample scRNA-seq data. It facilitates convenient exploration of multiple feature types, multivariate phenotype, and multiple datasets. It can also help an investigator to compare results from multiple datasets and data analysis protocols and evaluate and adjust for the impacts of potential confounders. It adds a new tool to the toolbox of computational biologists which we hope will eventually make the complicated analysis of multi-sample scRNA-seq data easily accessible to an average investigator to accelerate the discovery of cellular and molecular mechanisms behind various normal and disease phenotypes using such data.

## Methods

### TreeCorTreat pipeline

#### Input data

The input for TreeCorTreat consists of three components: a gene expression matrix *E*(*G*×*C*, with rows representing *G* genes and columns representing *C* cells), cell-level metadata *M*(*C*×*K*, where *K* ≥ 2) and sample-level metadata *L*(*S*×*J*, where *J* ≥ 2). To avoid redundancy and save storage space, we split metadata into cell-level and sample-level metadata. Cell-level metadata primarily annotates cell-level information and must contain at least 2 columns for each cell: cell barcode and corresponding sample identifier (ID). An optional third column can be added to provide cell-type annotation. Cell barcode is used to couple cell-level metadata *M* and gene expression matrix *E*. Sample-level metadata documents a sample’s phenotype(s) of interests (e.g. clinical outcome) and other related covariates (e.g. age and sex). The first column contains unique sample IDs, and the remaining columns contain phenotypes and covariates. The cell-level metadata and sample-level metadata can be linked via unique sample IDs.

#### Module 1 - Integration

By default, Harmony algorithm is applied to integrate cells from different samples and embed them into a common low-dimensional space [16]. This algorithm uses soft clustering to assign cells into distinct clusters and learns a cell-specific correction factor iteratively after projecting cells to a low-dimensional space (e.g. principal component space). We adapt the ‘*RunHarmony*’ and ‘*BuildClusterTree*’ functions from Seurat v3 [17] and wrap the following steps into a standalone ‘*treecor_harmony()*’ function with tunable parameters: library size normalization, integration feature selection, Principal Component Analysis (PCA), Harmony integration, unsupervised louvain clustering, Uniform Manifold Approximation and Projection (UMAP), hierarchical clustering of louvain clusters, and differentially expression analysis to identify cell type marker genes to facilitate annotation. Specifically, for library size normalization, gene counts for a given cell are divided by total counts for that cell, multiplied by a scaling factor (10^4^) and applied a natural log transformation. Features for integrating samples are obtained by choosing highly variable genes (HVGs) based on variance-mean relationship within each individual sample (using ‘*SelectIntegrationFeatures*’ function) and ranking the features based on the number of samples where they are identified as HVGs. By default, top 2000 HVGs are chosen and fed into downstream PCA procedure. Harmony is carried out based on the top 20 PCs and the corrected harmony embedding is obtained. Top 20 harmony coordinates are then used for downstream louvain clustering and UMAP analyses (using ‘*FindClusters*’ and ‘*RunUMAP*’ functions in Seurat). Instead of using this default integration pipeline, TreeCorTreat also allows users to provide their own integration parameters. Users can also use their own integration results in a Seurat object or cell cluster labels in an additional ‘celltype’ column in the cell-level metadata and skip the integration step. For hierarchical clustering of louvain clusters, ‘*BuildClusterTree*’ function is used. To facilitate cell type annotation, cell type specific genes are identified by comparing cells in each cluster versus cells in all other clusters via the Wilcoxon rank sum test using the ‘*FindAllMarkers*’ function in Seurat.

#### Module 2 - Define cell type hierarchical structure

The previous steps defined cell clusters either by louvain clustering or based on user-provided cell-level metadata (i.e. the optional column 3 in the cell metadata). Users can add text labels to annotate the cell type of each cell cluster based on top differentially expressed genes or examining known cell type marker genes. To facilitate association analyses at multiple resolutions, cell clusters are further clustered hierarchically. The hierarchical clustering tree is used to define cell types at coarser levels. The tree can be derived using either a data-driven approach or a knowledge-based approach. In the data-driven approach, the tree is generated by the hierarchical clustering of the louvain clusters in our ‘*treecor_harmony()*’ function. Specifically, PC scores are first averaged across cells within each cell cluster and a hierarchical tree is constructed (based on Euclidean distance and complete linkage method for hierarchical clustering). In the knowledge-based approach, users can specify the tree based on their prior knowledge by providing a string to describe the parent-children relationship of clusters at different granularity levels, as described in the Results section. The string utilizes nested parentheses and special characters to encode the basic tree structure. The leaf nodes in the tree correspond to cell clusters obtained from either the louvain clustering in the integration step or the cell metadata. In both data-driven and knowledge-based approaches, users can modify the cell type label via ‘*modify_label()*’ function.

#### Module 3 - Cell Type Proportion Analysis

Analyses of cell type proportions are wrapped into a standalone function ‘*treecor_ctprop()*’. Here we first define the feature for each tree node and then evaluate the association between the feature and the sample phenotype.

For a leaf node cell cluster, the feature is defined as the proportion of cells in a sample that fall into that node. For an intermediate parent node or root node, users can choose to define the feature in one of the three different ways listed below.

‘*Aggregate*’ (default setting): All cells from its descendant nodes are treated as a combined cell cluster. The feature of the node is defined as the proportion of cells in a sample that fall into this combined cluster. This is a univariate feature.
‘*Concatenate leaves’:* All terminal leaf nodes descending from the current node are collected, and their features (i.e. cell type proportions) are concatenated into a vector. The vector is the feature for the current node. This feature is a multivariate vector.
‘*Concatenate immediate children*’: All immediate children of the current node are collected. The feature for each immediate child is obtained via ‘*Aggregate*’ which gives the cell type proportion of that immediate child node. The features from all immediate children are then concatenated into a vector to serve as the feature for the current node. This feature is a multivariate vector.

Once the feature for each node is defined, we will go through each tree node to examine the correlation between its feature and the sample phenotype. The way to compute the correlation depends on whether the phenotype is univariate or multivariate.

*Univariate phenotype:* When the cell type proportion feature is univariate, the Pearson (default) or Spearman correlation between the feature and the phenotype across samples is computed. One can also choose a regression approach, where cell proportion serves as a response variable and the phenotype serves as an explanatory variable. A likelihood ratio test is performed comparing a full model with the phenotype included in the explanatory variables and a reduced model without the phenotype variable, and the likelihood ratio statistic is reported. When the cell type proportion feature is multivariate, by default the canonical correlation is computed. In other words, we find the optimal linear combination of feature variables that maximizes the correlation with the phenotype and report the maximized correlation. This optimization is directly solved by the canonical correlation analysis (CCA) [71]. In addition to CCA, one can also choose a regression approach where the cell proportion feature vector serves as response and the phenotype serves as an explanatory variable. Multivariate ANOVA (MANOVA, Pillai-Bartlett) is performed to compare two models with and without the phenotype in the explanatory variable, and a F-statistic is reported. Note that MANOVA can be viewed as a special case of regression.
*Multivariate phenotype analyzed separately:* One can analyze multivariate phenotype either separately or jointly. When the traits in the multivariate phenotype are analyzed separately, we treat each trait as a univariate phenotype and run the *univariate phenotype* analysis as above, but the results for all traits will be visualized in the same TreeCorTreat plot as in **Fig. 5A**.
*Multivariate phenotype analyzed jointly:* When the traits in the multivariate phenotype are analyzed jointly, by default the canonical correlation between the feature (either univariate or multivariate) and the multivariate phenotype is computed. In other words, CCA is used to find the optimal linear combination of feature variables and the optimal linear combination of phenotype variables that maximizes the correlation between the two. The canonical correlation is the maximized correlation. In addition to CCA, one can choose a regression approach where the cell type proportion feature serves as response and phenotypic traits as explanatory variables. For an univariate feature (Aggregate), likelihood ratio statistic is reported. For a multivariate feature (Concatenate), a MANOVA F-statistic is reported.

Note that for CCA, when the feature dimension or the phenotype dimension (i.e. the number of variables in the feature or phenotype) is larger than the sample size (i.e. the number of scRNA-seq samples), the problem is not full rank and one can trivially get canonical correlation=1 which is meaningless. In those situations, users should first reduce the dimension of the feature or phenotype before CCA. In order to make the results stable, when TreeCorTreat encounters a multivariate feature or phenotype in CCA, its default setting will always perform a dimension reduction using principal component analysis (PCA) first. The PCA will reduce the multivariate feature or phenotype vector to *k* dimensions *(k=2* by default), and then CCA will run using the data with the reduced dimension. Users have the option to set the dimension *k* to other values. Similarly, PCA is also used to reduce the dimension of multivariate features in the regression setting.

To determine the statistical significance of the association statistic (correlation, likelihood ratio statistic, or F-statistic), p-values are computed based on a permutation test. In other words, samples’ phenotype labels are shuffled multiple times (n=1000 by default), and the same procedure for computing the association statistic is run to construct a null distribution. For each node, the p-values are computed as the fraction of permutations with the association statistic as extreme as or more extreme than the observed one. Benjamini & Yekutieli (BY) procedure [58] is used to adjust p-values to account for multiple testing for dependent tests across all tree nodes. The adjusted p-value≤0.05 is used as the default significance cutoff, but users can also specify their own cutoffs.

A summary of the analysis procedures and corresponding R commands for different scenarios is provided in **Additional file 3: Table S3**.

#### Module 4 - Global Gene Expression Analysis

Analyses of global gene expression are wrapped into a standalone function ‘*treecor_expr()*’. Here we also first define the feature for each tree node and then evaluate the association between the feature and the sample phenotype.

For a leaf node, we first pool all cells in the node to obtain a pseudobulk profile of the node. Pseudobulk profiles are normalized by library size. We then use these pseudobulk profiles from all samples to select highly variable genes (HVGs) using locally weighted scatterplot smoothing (LOWESS) procedure in R. In other words, a gene-specific LOWESS function of standard deviation against its mean is fitted. By default, genes with positive residuals will be selected as HVGs. Users also have an option to specify how many HVGs to be selected. The pseudobulk profile of the selected highly variable genes in each sample is used as the feature vector of the node. This feature vector is multivariate.

For an intermediate parent node or root node, users can choose to define the feature in one of the three different ways listed below.

‘*Aggregate*’ (default setting): All cells from its descendant nodes are treated as a combined cell cluster. Cells in the cluster are pooled to create an aggregated pseudobulk profile. Pseudobulk profiles are normalized by library size. We use these pseudobulk profiles from all samples to select highly variable genes. The pseudobulk profile of the selected highly variable genes in each sample is used as the feature vector of the current node. This feature vector is multivariate.
‘*Concatenate leaves*’: All terminal leaf nodes descending from the current node are collected, and their feature vectors (i.e. gene expression vectors of highly variable genes) are concatenated into one vector to serve as the feature vector for the current node. This feature vector is multivariate.
‘*Concatenate immediate children*’: All immediate children of the current node are collected. The feature vector for each immediate child is obtained via ‘*Aggregate*’ which gives the pseudobulk profile of highly variable genes of that immediate child node. The features from all immediate children are then concatenated into one vector to serve as the feature vector for the current node. This feature vector is multivariate.

Once the feature vector for each node is defined, we will go through each tree node to examine the correlation between its feature and the sample phenotype as below.

*Univariate phenotype:* PCA is used to reduce the dimension of the feature vector to *k* (*k*=2 by default). The canonical correlation between the phenotype and the dimension reduced feature vector is calculated using CCA and reported. Optionally, one can also choose a regression approach, where the top *k* PCs of the feature serve as the response and the sample phenotype serves as an explanatory variable. Multivariate ANOVA (MANOVA) is performed to compare two models with and without the phenotype in the explanatory variable, and a F-statistic is reported.
*Multivariate phenotype analyzed separately:* When the traits in the multivariate phenotype are analyzed separately, we treat each trait as a univariate phenotype and run the Univariate phenotype analysis as above. The results for all traits will be visualized in the same TreeCorTreat plot similar to **Fig.5A**.
*Multivariate phenotype analyzed jointly*: When the traits in the multivariate phenotype are analyzed jointly, by default the canonical correlation between the first *k* PCs (*k*=2 by default) of the feature vector and the multivariate phenotype is computed using CCA and reported. When the phenotype dimension is larger than the sample size, we will also first reduce its dimension by PCA (default *k*=2) before running CCA. Optionally, one can also choose a regression approach, where the top *k* PCs of the feature serve as the response and the phenotypic traits serve as explanatory variables. Multivariate ANOVA (MANOVA) is performed to compare two models with and without the phenotype in the explanatory variable, and a F-statistic is reported.

Similar to the analysis of cell proportions, the statistical significance of each node is determined by p-values computed using a permutation test, where the null distribution of association statistic (correlation or F-statistic) is constructed by shuffling samples’ phenotype labels multiple times (n=1000 by default) and then running the same procedure for computing the association statistic. The p-values are then adjusted for multiple testing using the BY procedure. The adjusted p-value ≤ 0.05 is used as the default significance cutoff, but users can also specify their own cutoffs.

#### Module 5 - Phenotype Associated Differential Expression Analysis

Analyses of differential gene expression are wrapped into a standalone function ‘*treecor_deg()*’. Here we first compute the pseudobulk gene expression profile for each tree node using the ‘*Aggregate*’ approach as before. In other words, for each node all cells from its descendants are pooled and aggregated into a pseudobulk profile in each sample. Pseudobulk profiles are normalized by library size. Limma [53] was then used to conduct differential expression analysis for each node as follows.

*Univariate phenotype:* An limma model was fit to identify phenotype associated differentially expressed genes (DEGs) by regressing gene expression against phenotype (e.g. healthy donors versus severe COVID-19 patients) with covariate adjustment (e.g. age). DEGs passing a user-specified false discovery rate (FDR) cutoff (default cutoff = 0.05) will be reported for each node and saved into csv files along with log fold change, t-statistics, p-value and FDR. The number of DEGs for each node is shown as a TreeCorTreat plot.
*Multivariate phenotype analyzed separately:* When the traits in the multivariate phenotype are analyzed separately, we treat each trait as a univariate phenotype and run the *univariate phenotype* analysis as above. The results for all traits will be visualized in the same TreeCorTreat plot similar to **Fig. 5A**.
*Multivariate phenotype analyzed jointly:* Here multiple traits are first combined into one variable either using a data-driven approach or user-specified weights. In the data-driven approach, the first principal component of the trait vector will be analyzed as a univariate phenotype. In the user-specified weight approach, traits will be aggregated into a univariate phenotype via a linear combination defined by user-provided weights. The aggregated trait will be analyzed as a univariate phenotype.

#### Module 6 - Visualization by TreeCorTreat plot

Users can flexibly visualize the results to pinpoint important cell types by plotting the decorated tree, using different colors, sizes or transparencies. Multiple plotting features are provided, such as tree skeleton representation (e.g. straight line, classical angle bend and quadratic bezier curve) and advanced data visualization (e.g. balloon plot, barplot and heatmap). More advanced plotting features have been demonstrated in **Fig. 3** and in the online user manual.

#### Adjusting for covariates

For computing Pearson, Spearman, or canonical correlation, a regression model can be used to regress cell type proportion or global gene expression profile (sample-level pseudobulk) against covariates (e.g. potential confounders) that users want to adjust for. Residuals will be treated as adjusted cell type proportion or gene expression in the correlation analysis. When phenotype-cell type association is analyzed using regression or MANOVA type methods, covariates can be adjusted directly by including them in the regression models.

### Data collection

For COVID-19 PBMC studies, processed read counts from Schulte-Schrepping et al. and Su et al. were downloaded from the European Genome-Phenome Archive (EGA) and ArrayExpress database using the accession number provided in the original publication: EGAS00001004571 and E-MTAB-9357. **Additional file 4: Table S4** contains sample-level metadata. In the sample-level metadata, demographics information such as age was obtained from the original papers, and disease severity of patients was either described in the original papers or inferred based on WHO ordinal score (WOS; mild: WOS=1-2; moderate: WOS=3-4; severe: WOS=5-7). For the NSCLC study, processed read counts were downloaded from the GEO under accession number GSE176021. Sample-level metadata are extracted from the original publication. All processed data were aligned to the GRCh38 genome.

### Data preprocessing

For the COVID-19 analysis, only PBMC samples were kept for subsequent analyses and thus 11 whole blood samples were excluded. Genes detected across all 3 COVID-19 cohorts were retained. Cells from each sample were filtered according to the following criteria: (1) the number of detected features (i.e. genes with positive count) per cell should be at least 500; (2) the proportion of mitochondrial gene counts must be less than 20%. After filtering cells, samples with less than 500 cells were further eliminated and genes with zero expression across all cells were removed. In total, 25,426 genes, 335 samples (i.e. 27 Healthy, 135 Mild, 76 Moderate and 97 Severe) and 659,375 cells were retained for downstream analyses.

For the NSCLC analysis, the preprocessing was done using the preprocessing script provided by the original publication [43]. Briefly, the script includes the following steps. Low-quality cells with >10% mitochondrial gene counts or <10% ribosomal gene counts or an outlying number of detected features were filtered out. A total of 560,916 T cells were retained. Dissociation associated genes, mitochondrial genes (starting with ‘MT-’), high abundance lincRNA genes, genes linked with poorly supported transcriptional models (with a prefix of ‘RP-’) and T cell receptor (TCR) genes (TRA/TRB/TRD/TRG) were excluded for subsequent analysis. Cells from 31 samples were integrated using Seurat MultiCCA (Seurat v3). The top 3,000 HVGs were chosen as features for integration. Genes associated with type I interferon response (IFN), immunoglobulin genes and mitochondrial related genes were additionally removed from clustering. Cells were clustered using louvain clustering, and cell clusters were annotated based on canonical immune cell markers and top differentially expressed genes within each cluster identified by Wilcoxon rank sum test.

## Declarations

## Acknowledgements

We thank Jiajia Zhang for sharing the codes for NSCLC data preprocessing.

## Availability of data and materials

The data used in this analysis are all publicly available. COVID-19 PBMC single-cell RNA-seq data can be downloaded from the European Genome-Phenome Archive (EGA) and ArrayExpress database using the accession number: EGAS00001004571 and E-MTAB-9357. NSCLC dataset can be downloaded from the Gene Expression Omnibus under the accession number GSE176021. TreeCorTreat is freely available as an open source R package under the GNU General Public License (version 2). The software and source code can be accessed from the GitHub repository (https://github.com/byzhang23/TreeCorTreat).

## Contributions

HJ conceived the study. BZ and HJ developed methods. BZ implemented the methods and developed the software. BZ and ZJ processed scRNA-seq data. BZ and HJ interpreted the results. BZ and HJ wrote the manuscript. All authors read, revised and approved the final manuscript.

## Ethics approval and consent to participate

Not applicable.

## Consent for publication

Not applicable.

