## Additional file 5: Supplementary Figures for "Tree-based Correlation Screen and Visualization for Exploring Phenotype-Cell Type Association in Multiple Sample Single-Cell RNA-Sequencing Experiments"

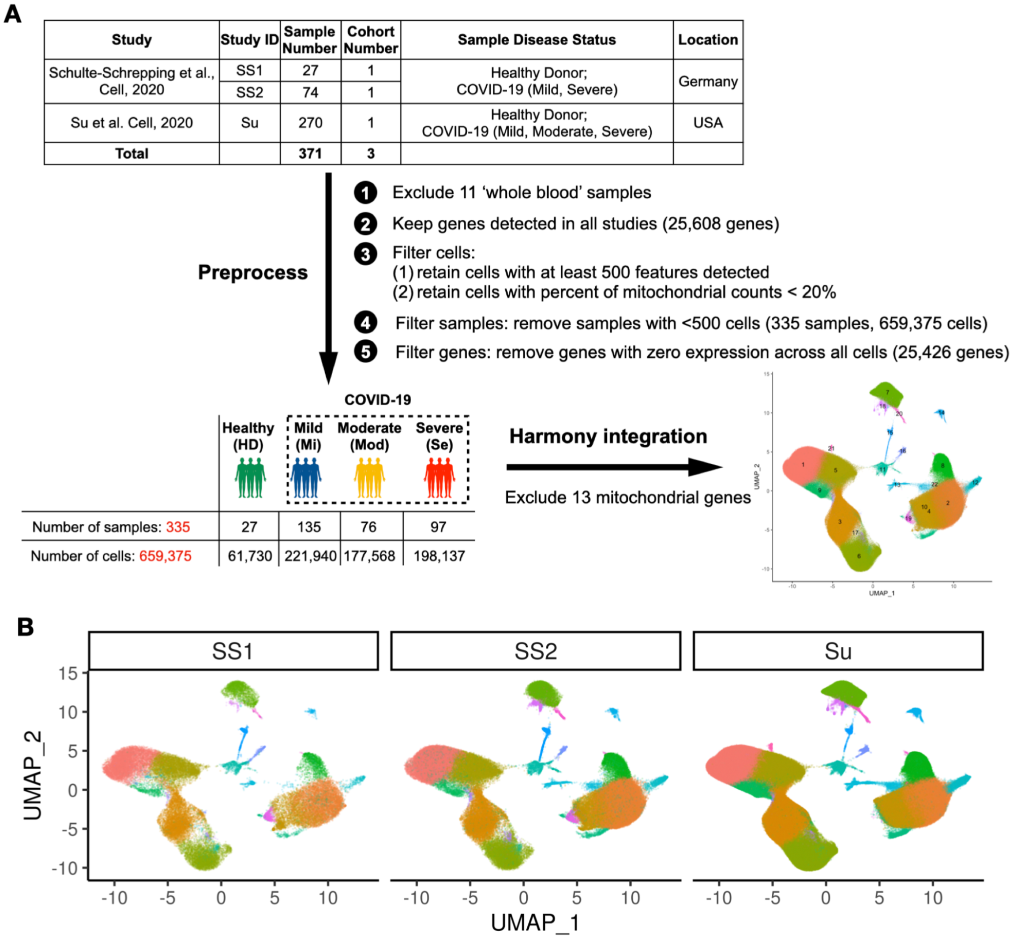


| Study | Study ID | Sample Number | Cohort Number | Sample Disease Status | Location |
|---|---|---|---|---|---|
| Schulte-Schrepping et al., Cell, 2020 | SS1 | 27 | 1 | Healthy Donor; COVID-19 (Mild, Severe) | Germany |
| | SS2 | 74 | 1 | | |
| Su et al. Cell, 2020 | Su | 270 | 1 | Healthy Donor; COVID-19 (Mild, Moderate, Severe) | USA |
| **Total** | | **371** | **3** | | |

**Figure S1.** (A) Preprocessing process of COVID-19 datasets. (B) UMAP stratified by three cohorts (SS1, SS2 and Su).

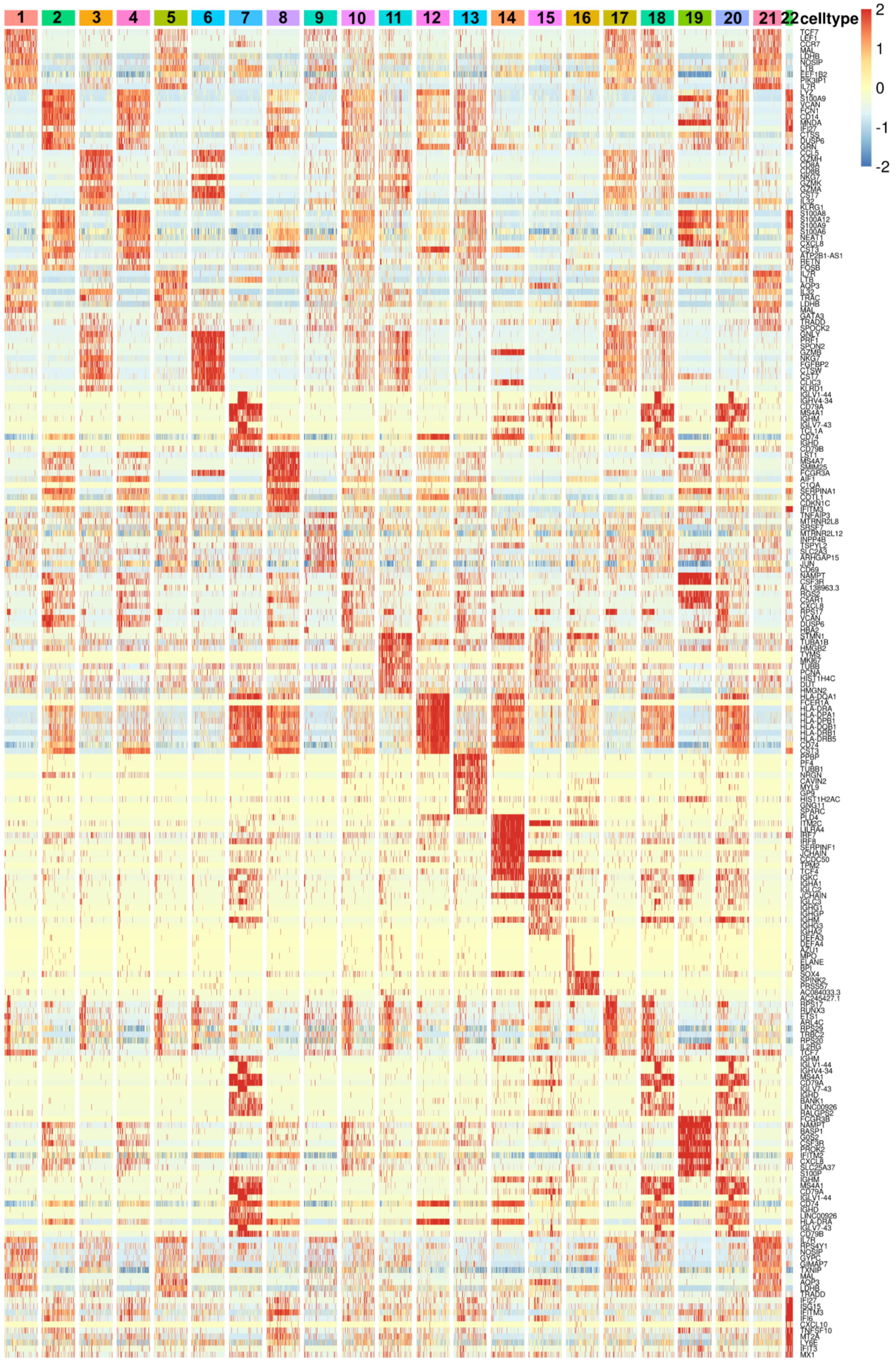


**Figure S2.** Top 10 differentially expressed genes for identified cell types. Five-hundred cells (or all cells in a cluster if that cluster had less than 500 cells) were randomly sampled from each cluster for visualization. Cell types were annotated by cluster ID.

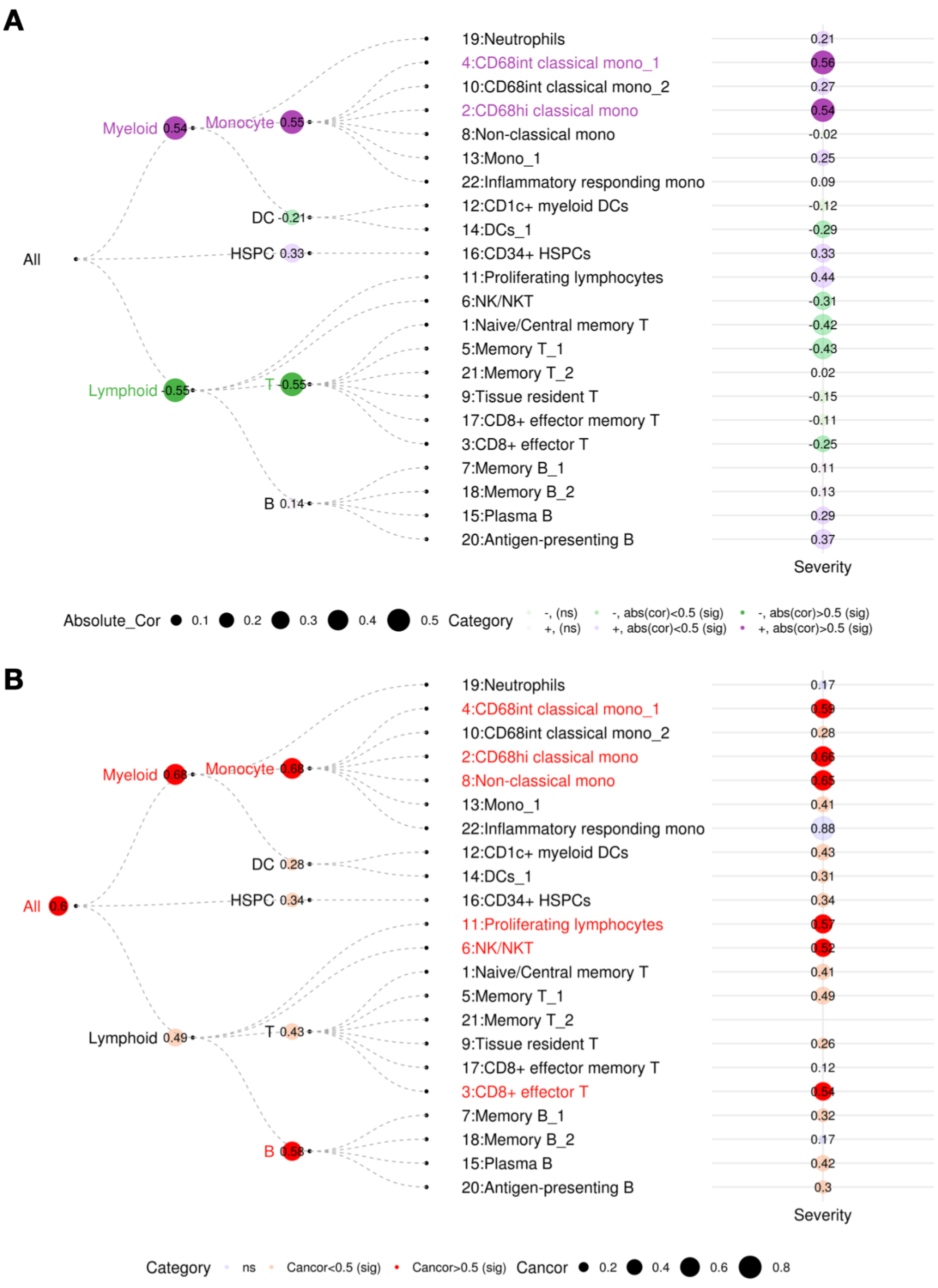


**Figure S3. Examples of TreeCorTreat plot with annotated numbers, using a single COVID-19 dataset (Su).** (A) TreeCorTreat plot of cell type proportion-diseases severity association, same as Fig. 2B except that the plot here is annotated with Pearson correlation for each cell type. (B) TreeCorTreat plot of global gene expression-diseases severity association, same as Fig.2D except that the plot here is annotated with canonical correlation annotation for each cell type.

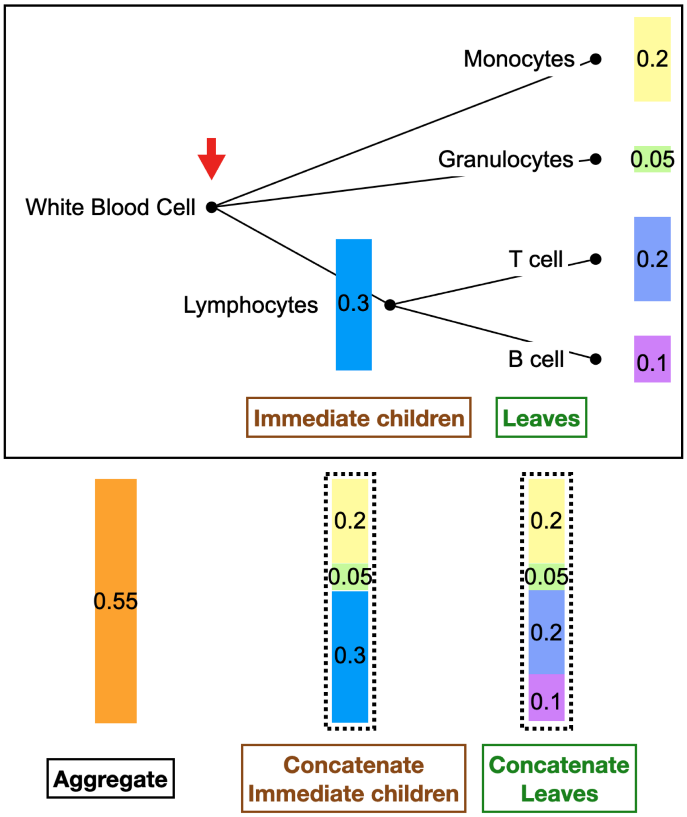


**Figure S4. A cartoon illustration of three feature summary approaches.**

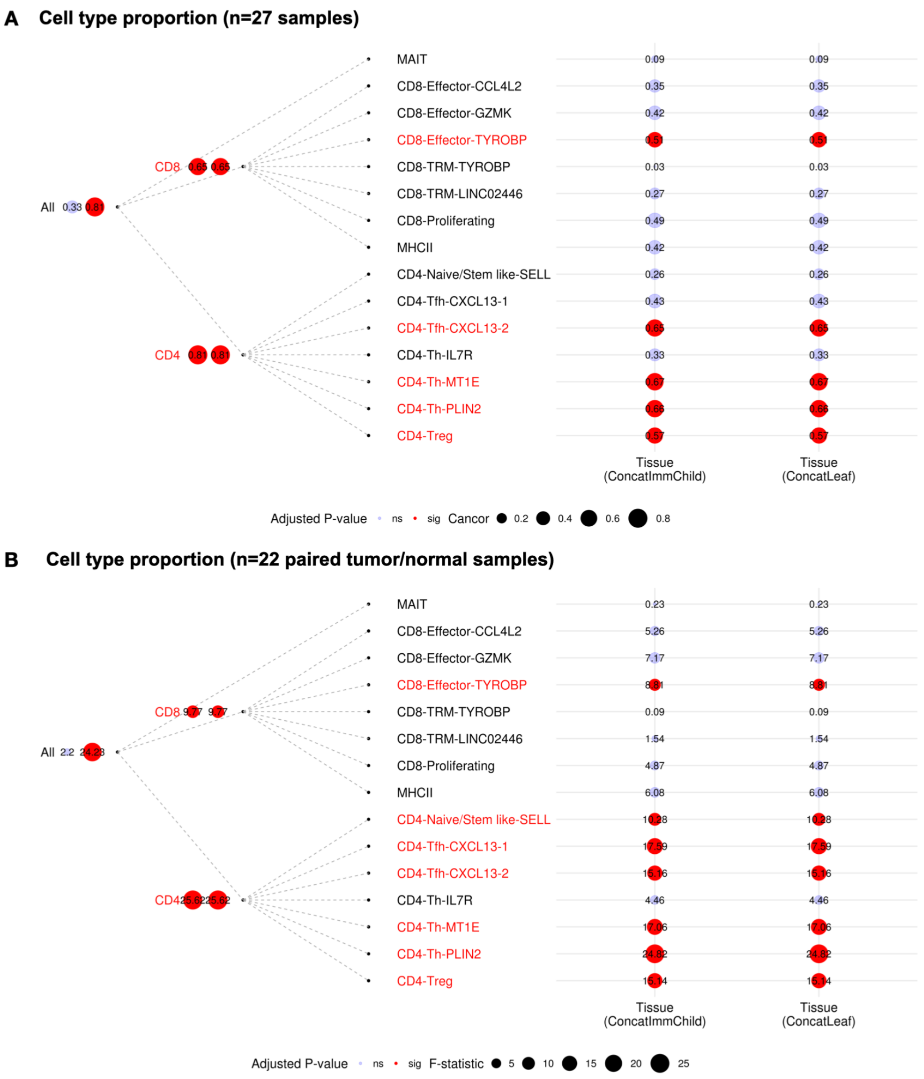


**Figure S5. TreeCorTreat plot of association between cell type proportion and tissue type in NSCLC data.** Results from two concatenation approaches in TreeCorTreat analysis. First column corresponds to 'Concatenate immediate children (ConcatImmChild)' and the second column corresponds to 'Concatenate leaf nodes (ConcatLeaf)'. Canonical correlation (instead of Pearson correlation) is shown for both leaf and internal nodes. Color indicates three categories defined by statistical significance: blue for not significant (ns) and red for adjusted p-value≤0.05 (sig). (A) Using 27 unpaired samples (15 TILs and 12 normal lungs) with circle size representing the magnitude of canonical correlation. (B) Using 22 paired samples (11 matching TIL-normal lung pairs) with circle size representing F-statistic. P-value was calculated based on 10,000 permutations and the BY procedure was used for multiple testing correction. Red colored labels highlight cell types with significant association with tissue compartments in both approaches (adjusted p-value≤0.05).

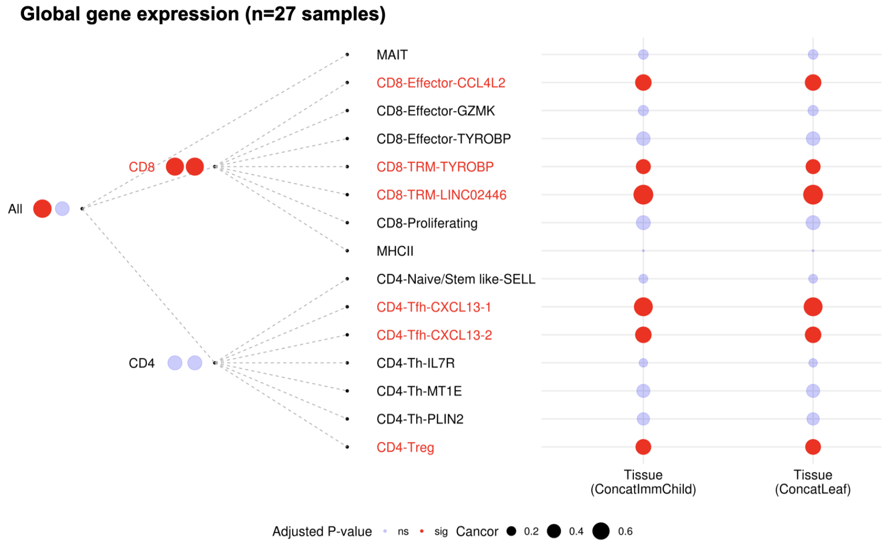


**Figure S6. TreeCorTreat plot of association between global gene expression and tissue type (n=27 samples; 15 tumor and 12 normal samples) in NSCLC data.** Results from two concatenation approaches in TreeCorTreat analysis. First column corresponds to 'Concatenate immediate children (ConcatImmChild)' and the second column corresponds to 'Concatenate leaf nodes (ConcatLeaf)'. Circle size represents the magnitude of canonical correlation and color indicates three categories defined by statistical significance: blue for non-significant (ns) and red for adjusted p-value≤0.05 (sig). P-value was calculated based on 10,000 permutations and the BY procedure was used for multiple testing correction. Red colored labels highlight cell types with significant association with tissue compartments in both approaches (adjusted p-value≤0.05).
